# Integrating Molecular QTL Data into Genome-wide Genetic Association Analysis: Probabilistic Assessment of Enrichment and Colocalization

**DOI:** 10.1101/078667

**Authors:** Xiaoquan Wen, Roger Pique-Regi, Francesca Luca

## Abstract

We propose a novel statistical framework for integrating genetic data from molecular quantitative trait loci (QTL) mapping into genome-wide genetic association analysis of complex traits, with the primary objectives of quantitatively assessing the enrichment of the molecular QTLs in complex trait-associated genetic variants and the colocalizations of the two types of association signals. We introduce a natural Bayesian hierarchical model that treats the latent association status of molecular QTLs as SNP-level annotations for candidate SNPs for complex traits. We detail a computational procedure to seamlessly perform enrichment, fine-mapping and colocalization analyses, which is a distinct feature compared to the existing colocalization analysis procedures in the literature. The proposed approach is computationally efficient and requires only summary-level statistics. We evaluate and demonstrate the proposed computational approach through extensive simulation studies and the analysis of blood lipid data and the whole blood eQTL data from the GTEx project. In addition, a useful utility from our proposed method enables the computation of expected colocalization signals, which is analogous to the power calculation in genetic association studies. Using this utility, we further illustrate the importance of enrichment analysis on the ability of discovering colocalized signals and the potential limitations of currently available molecular QTL data.

## 1 Introduction

Genome-wide association studies (GWAS) have successfully identified many genomic loci that impact complex diseases and complex traits. Nevertheless, the molecular pathways that connect genetic variants to complex traits are still poorly understood, largely because a considerable proportion of trait-associated signals are located in the non-coding region of the genome. With recent advancements in high-throughput sequencing technology, systematic investigations of cellular phenotypes have revealed an abundance of non-coding molecular quantitative trait loci (QTLs) (Ardlie *et al.*, 2015, McVicker *et al.*, 2013, Banovich *et al.*, 2014, Degner *et al.*, 2012). Integrating molecular QTL data into GWAS analyses has shown great potential in unveiling the missing links between trait-associated genetic variants and organismal phenotypes (Gamazon *et al.*, 2015, Nica *et al.*, 2010, Teslovich *et al.*, 2010).

In this paper, we focus on a specific type of integrative analysis that aims to assess the overlapping/colocalization of causal GWAS hits and causal molecular QTLs (also known as quantitative trait nucleotides, or QTNs). Following Giambartolomei *et al.* (2014), we define that a GWAS hit and a molecular QTN are colocalized if a single genetic variant is causally associated with both the complex and molecular traits of interest. Colocalizing genetic variants that jointly affect both molecular and organismal phenotypes provides an intuitive starting point for exploring the role of genetic variants in disease etiology. Taking expression quantitative trait loci (eQTL) mapping as an example, colocalizing an eQTL signal with a GWAS hit naturally suggests that the target gene of the eQTL may play an important role in the molecular pathway of the complex traits. Additionally, other types of available integrative analysis approaches, e.g., *Sherlock* (He *et al.*, 2013), *PrediXcan* (Gamazon *et al.*, 2015) and other similar approaches (Gusev *et al.*, 2016, Zhu *et al.*, 2016), can also benefit from accurate colocalization analysis, either for improved power (as in the case of *Sherlock*) or better interpretation of the inference results (as in the case of *PrediXcan*).

Considering a practical setting in which GWAS and molecular QTL data are obtained from different sets of samples, we propose a natural Bayesian hierarchical model for integrating the two types of association data. Specifically, we regard the (latent) association status of each candidate SNP with respect to the molecular phenotype of interest as an SNP-level annotation, and we attempt to quantify the odds of an annotated SNP being causally associated with the complex trait of interest, which is statistically equivalent to evaluating the enrichment level of annotated SNPs in the causal GWAS hits. Subsequently, the resulting enrichment estimates are utilized in the downstream fine-mapping (of GWAS hits) and colocalization analyses. Within our Bayesian hierarchical model, we show that the problems of enrichment estimation, fine-mapping and colocalization testing can be seamlessly solved in a unified inference framework. In addition, our approach is computationally efficient and requires only summary-level data from both molecular QTL mapping and GWAS.

Our proposed method is most similar to the probabilistic model-based approaches *coloc* (Giambartolomei *et al.*, 2014) and *eCAVIAR* (Hormozdiari *et al.*, 2016), which represent the state-of-the-art in the current literature. The advantages of the model-based colocalization analysis methods over the empirical methodologies (e.g., Nica *et al.* (2010)) have been fully demonstrated through both rigorous theoretical arguments (Wallace, 2013, Giambartolomei *et al.*, 2014) and carefully constructed simulation studies (Hormozdiari *et al.*, 2016). In this paper, we show that both *coloc* and *eCAVIAR* can be viewed as special cases of the proposed approach with additional simplifying assumptions. Consequently, our approach shares the advantages of both existing approaches, but it enjoys additional flexibility and improved statistical rigor.

## 2 Method

### 2.1 Model and Notation

Without loss of generality, we consider a GWAS of a quantitative trait and describe its associations with *p* candidate SNPs and *n* unrelated samples using a multiple linear regression model,

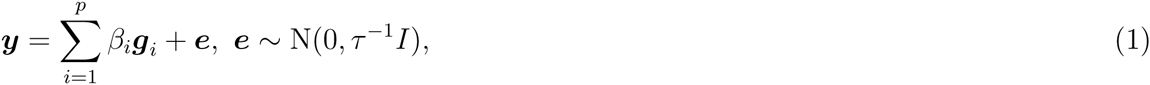

where we assume that both the phenotype and genotypes are centered (the intercept term is therefore exactly 0) and denote the complete collection of genotypes as ***G*** := [***g***_1_,…, ***g***_*p*_]. We further denote the latent binary association status of each SNP *i* by dichotomizing its genetic effect *β*_*i*_, i.e., *γ*_*i*_ = 1 indicates that SNP *i* is causally associated and *β*_*i*_ ≠ 0 otherwise. It can be argued that the aim of the GWAS is to make inference of the binary vector ***γ*** := (*γ*_1_,…, *γ*_*p*_). In addition, we assign the standard spike-and-slab prior for each regression coefficient *β*_*i*_ and a flat gamma prior for the residual error variance parameter *τ*.

Suppose that a quantitative annotation (categorical or continuous) is available for each candidate genetic variant. We integrate the SNP-level annotation into the association analysis by specifying a natural logistic prior for each candidate SNP *i*, i.e.,

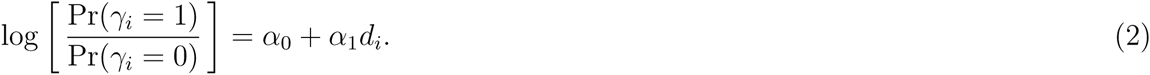

In particular, we denote the complete collection of the SNP annotation data as ***d*** := (*d*_1_,…, *d*_*p*_), and we refer to ***α*** := (*α*_0_, *α*_1_) as the enrichment parameter: for a binary annotation, a positive *α*_1_ value indicates that SNPs with the feature have increased odds of being associated with the trait of interest, i.e., the annotated feature is enriched in the trait-associated genetic variants.

In this paper, we consider a special setting in which the annotation is derived from the association analysis of molecular QTL data, namely, (***Y***_*qtl*_, ***G***_*qtl*_). Intuitively, the true association status of each SNP with the molecular phenotype can be naturally incorporated as annotations for GWAS analysis. However, due to the intrinsic limitations in the molecular QTL mapping, e.g., imperfect power and complication of LD among SNPs, the desired binary association status of each SNP with respect to the molecular phenotype of interest, ***d***, is practically impossible to obtain. Rather, we propose using the posterior distribution of ***d***, Pr(***d*** | ***Y*** _*qtl*_, ***G***_*qtl*_), to represent a “fuzzy” version of the annotation to naturally account for the uncertainty. Specifically, we modify the prior model (2) and regard the annotation data ***d*** as unobserved but a realization from the *joint* posterior distribution obtained from the Bayesian multi-SNP association analysis of the molecular QTL data, i.e.,

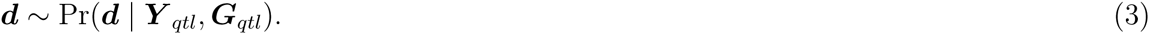

Furthermore, the uncertainties attached to *both* ***γ*** and ***d*** from the genetic association analysis give rise to the problem of colocalization: it is natural to quantify the colocalization of association signals for SNP *i* using the following probability statement

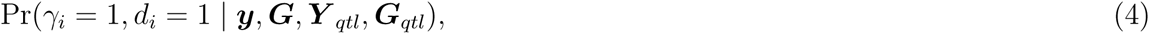

where both *γ*_*i*_ and *d*_*i*_ are regarded as random variables.

Given observed complex trait data, (***y***, ***G***), and the true annotation ***d*** (assuming observed), our recent work (Wen *et al.*, 2016) demonstrated that the proposed Bayesian hierarchical model can be efficiently fit based on an algorithm named deterministic approximation of posteriors (DAP) by adopting an empirical Bayes strategy. In brief, the inference procedure divides the genome into roughly independent LD blocks (Berisa and Pickrell, 2016) and proceeds in two stages of analysis. In the first stage, we obtain the maximum likelihood estimate of the enrichment parameter, 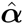, using an EM algorithm by treating ***γ*** as missing data. Subsequently, in the second stage, we perform multi-SNP fine-mapping conditional on 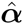 within each LD block and report the posterior distribution of ***γ***. In the following sections, we discuss an extended inference framework to account for the fuzzy annotations in the prior model (3) and solve the colocalization problem.

### 2.2 Enrichment Analysis of Molecular QTLs in GWAS Hits

The primary objective of the enrichment analysis is to estimate the hyper-parameter ***α*** given the observed summary statistics from GWAS and the fuzzy annotation of molecular QTLs. Note that if the binary molecular QTL annotation is indeed known, then the EM algorithm described in Wen *et al.* (2015, 2016) can be directly applied to obtain the maximum likelihood estimate of ***α***. With incomplete information on annotation data, we adopt a principled statistical strategy in missing data inference known as *multiple imputation* (Rubin, 1987, Little and Rubin, 2002). Specifically, the multiple imputation procedure creates *m* complete data sets by filling in, i.e., imputing, the missing entries of the binary annotation data. The imputed data sets are then individually analyzed using the existing EM algorithm, and the distinct estimates of 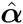 from multiple imputed data sets are combined into a final estimate using a set of rather simple rules (Appendix A). The key to implementing this strategy is to impute the annotations, which, in our case, is achieved by sampling from the posterior distribution Pr(***d*** | ***Y*** _*qtl*_, ***G*** _*qtl*_). Note that sampling from the *joint* posterior distribution (compared to simply using the marginal posterior inclusion probability of each SNP) preserves the uncertainty from identifying causal molecular QTNs due to LD, particularly in the presence of multiple *cis*-eQTL signals, and improves the accuracy of the enrichment estimate.

The number of imputed data sets (*m*) necessary for reliable estimation has been systematically studied in the missing data theory. The common consensus in the statistical literature is that *m* should be determined by the percentage of missingness, and various theoretical and empirical studies (Schafer, 1999, Graham *et al.*, 2007) roughly agree that 20 imputations are required for 10% to 30% missing information and that 40 imputations are required for 50% missing information. Although the true annotation ***d*** is completely unobserved in our context, we are certain that *d*_*i*_ = 0 for the vast majority of the candidate SNPs by inspecting the posterior distribution Pr(***d*** | ***Y***_*qtl*_, ***G***_*qtl*_). In fact, by examining the analysis results of *cis*-eQTLs from the GTEx whole blood data, we find that there are only *∼* 1.5% *cis* candidate SNPs with a posterior inclusion probability *≥* 0.01. Guided by this empirical evidence, we choose to impute *m* = 25 QTL data sets for each analysis. (We have also experimented with 50 and more imputed data sets in the simulations, and the inference results are virtually unchanged.)

Additionally, we observed that detectable GWAS hits and eQTLs (with currently available sample sizes) are both relatively sparse in practice, which can lead to large variances for the estimated enrichment parameter *α*_1_. To illustrate this point, we consider that both ***γ*** and ***d*** are observed; it is then trivial to estimate 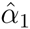 using a 2 × 2 contingency table. Because each binary vector contains only very few non-zero entries, the resulting contingency table is extremely imbalanced. Consequently, the variance of 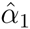 (approximately equal to the inverse of the smallest cell count) can be large, and the point estimate can be unstable. To stabilize the estimate of the enrichment parameter, we modify the original EM algorithm and apply an *l*_2_ penalty with a shrinkage parameter *λ* in the M-step to shrink the estimate toward 0. This strategy is informed by the statistical principle of “variance-bias trade-off”. Alternatively, this can be viewed as assigning a N(0, 1*/λ*) prior to *α*_1_. In practice, we select *λ* in a data-driven manner by assessing the degree of imbalance of the unobserved contingency table based on the association data (Appendix B), which assigns stronger penalties for larger degrees of imbalance.

### 2.3 Fine-mapping Incorporating Molecular QTL Annotations

Given the point estimate of the enrichment parameter, we adopt an empirical Bayes procedure to infer the true association status, *γ*, for all SNPs in GWAS. Specifically, the prior for each SNP *i* in multi-SNP fine-mapping can be computed in a straightforward manner by a two-component mixture, i.e.,

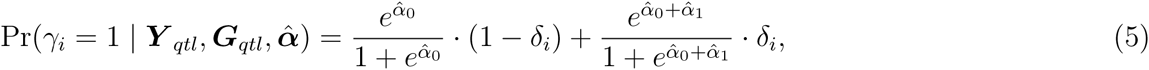

where *δ*_*i*_ := Pr(*d*_*i*_ = 1 | ***Y*** _*qtl*_, ***G*** _*qtl*_), i.e., the marginal posterior inclusion probability (PIP) of SNP *i* being a causal molecular QTN obtained from the eQTL mapping.

Given the prior (5), the adaptive DAP algorithm described in (Wen *et al.*, 2016) can be applied to each LD block separately for multi-SNP fine-mapping. The inference results are represented by the joint distribution of ***γ*** and by the marginal posterior inclusion probability, 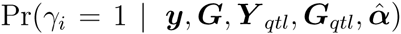, which highlights the importance of each individual SNP while accounting for LD. As with the enrichment estimate, in Appendix E, we show that the multi-SNP fine-mapping can be performed relying only on summary-level statistics from GWAS using the adaptive DAP algorithm.

Because our primary objective in this paper is to assess the colocalization signals (rather than identify multiple independent GWAS signals), we adopt an alternative strategy that efficiently computes approximate GWAS PIPs for putative associations of interest for colocalization analysis. For each pre-identified GWAS signal (from either single or multiple SNP association analysis), we specify an LD block that contains a set of candidate SNPs. Within each candidate SNP set, we assume that there exists at most a single causal GWAS hit and apply the DAP-1 algorithm using the summary-level statistics to perform fine-mapping analysis incorporating the prior model (5). Overall, this strategy is similar to the fine-mapping approaches described in Pickrell (2014), Veyrieras *et al.* (2008), Maller *et al.* (2012). However, in contrast to the aforementioned approaches, our prior specification allows the colocalization/fine-mapping analysis to utilize different LD block units that are completely independent of the estimation of the enrichment parameters. We call this strategy the “signal-centric” approach and demonstrate its use in our simulation and real data examples.

### 2.4 Colocalization Analysis of GWAS and Molecular QTL Data

Colocalization analysis aims to quantify the overlap between the *causal* GWAS hits and molecular QTNs at the SNP level. Within our probabilistic framework, it can be conveniently formulated as the evaluation of the joint posterior probability

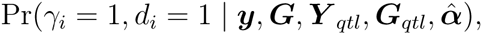

for each SNP *i*.

Given the fine-mapping PIP result 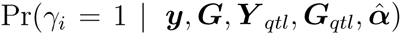, the colocalization probability for SNP *i* can be computed as

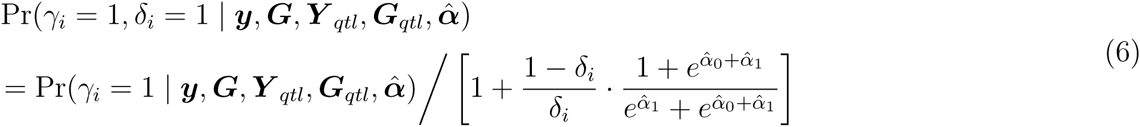

by solving a simple linear system (Appendix C). We refer to this quantity as the *SNP-level colocalization probability* (henceforth referred to as SCP).

Note that the resulting SNP-level colocalization probabilities still carry uncertainties due to LD. We demonstrate this point by considering a hypothetical example of two perfectly correlated SNPs and assuming that they are in complete linkage equilibrium with the remaining candidate SNPs. If one of the two SNPs is genuinely associated with the molecular phenotype, it should follow that *δ*_1_ = *δ*_2_ = 0.5 in a well-powered molecular QTL analysis, indicating the certainty that one of the SNPs is the causal QTN; however, there is no further information to distinguish the two. Consequently, the two SNPs carry the same prior for the GWAS analysis and obtain the same marginal posterior inclusion probabilities. In the case that one of the two SNPs is also genuinely associated with the complex trait (it is possible that the causal GWAS SNP is different than the molecular QTN), we should similarly find that 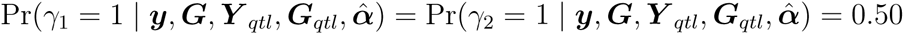 with sufficient power. Furthermore, as shown in Equation (6), the SNP-level colocalization probabilities for the two SNPs are also identical, with the actual value depending on the estimate of the enrichment parameter 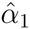: as 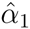 → 0, both take a value of 0.25, whereas when 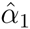 → ∞, both take a value of 0.50.

Following Guan and Stephens (2011), Wen *et al.* (2015), we propose computing a *regional colocalization probability*, or RCP, by summing up the SNP-level colocalization probabilities (SCPs) of correlated SNPs within an LD block. RCP is naturally interpreted as the probability of a genomic region harboring a colocalized signal. In our hypothetical example of two perfectly linked SNPs, the RCP → 1 if 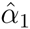 → ∞, which indicates near certainty that one of the SNPs is both a QTN and the causal GWAS hit (i.e., a colocalized signal); in contrast, when 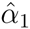 → 0, the resulting RCP value implies that there is an approximately 50% chance that the molecular QTL and the GWAS signals are *not* colocalized. We plot the functional relationship of RCP with respect to *α*_1_ in Figure 1, which illustrates the impact of the enrichment estimation on the quantitative assessment of colocalized signals. In general, we find that RCPs can be more informative and easier to interpret than SCPs in the presence of LD.

**Figure 1:**
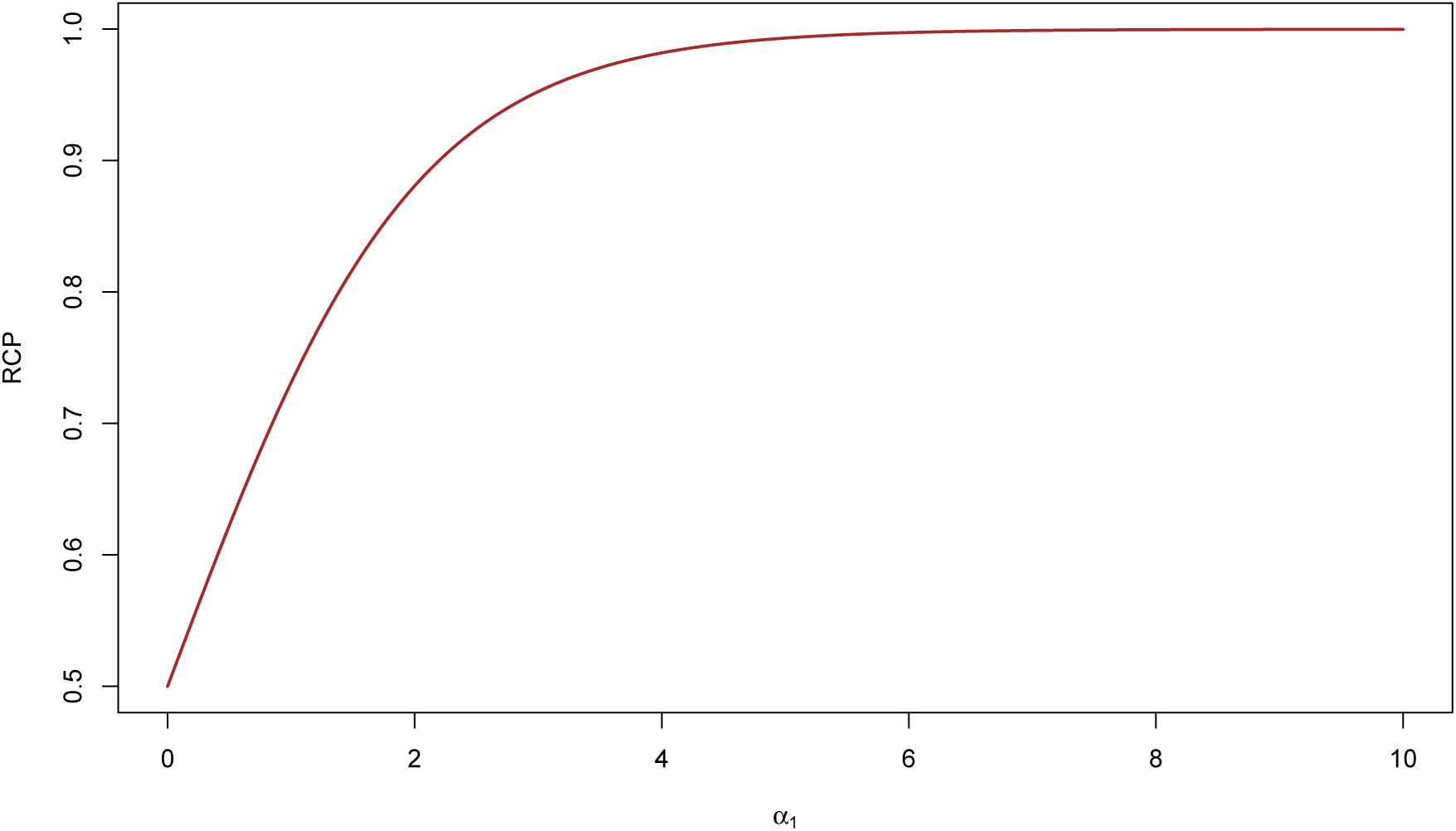
Functional relationship between RCP and enrichment parameter *α*_1_ in a hypothetical example. We consider two perfectly linked SNPs: one is causally associated with the molecular phenotype of interest, and one is causally associated with the complex trait of interest. Assuming that the two SNPs are in complete linkage equilibrium with other SNPs, the plot shows the functional relationship of the RCP value with respect to the enrichment parameter. Note that we should conclude that the two association signals are colocalized (RCP *→* 1) only if the enrichment level is sufficiently high. It is also theoretically possible that the RCP ≤ 0.5 if the molecular eQTLs are depleted in the GWAS hits, i.e., *α*_1_ < 0.

#### 2.4.1 Connection to Existing Probabilistic Colocalization Approaches

In this section, we show that Equation (6) represents a generalization of existing probabilistic approaches for colocalization analysis, namely, *eCAVIAR* and *coloc*. If we assume that eQTLs and GWAS hits are independent *a priori*, i.e., *α*_1_ is restricted to 0, then the prior for each SNP in GWAS becomes irrelevant to the eQTL data, and Equation (6) can be subsequently simplified to

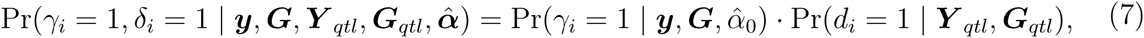

which coincides with the colocalization posterior probability (CLPP) proposed in *eCAVIAR*.

We present the detailed derivation that connects our general model to the *coloc* model in Appendix D and describe the additional simplifying assumptions made by *coloc*. More importantly, we show that the required prior probabilities in the *coloc* model can be equivalently parametrized by our enrichment parameter ***α***. However, in contrast to our proposed approach that estimates the enrichment parameter from the data, the prior probabilities in *coloc* are pre-specified subjectively.

#### 2.4.2 Bayesian Hypothesis Testing of Colocalization

In colocalization analysis, it is occasionally of interest to test the following hypothesis:

*H*_0_: Genomic region *i* does not contain a colocalized signal, for each locus *i*. Here, we show that the above hypothesis testing problem can be conveniently solved through the posterior inference within the proposed Bayesian framework.

Given a set of rejected hypotheses, *M*, the Bayesian false discovery rate (FDR) can be intuitively estimated by

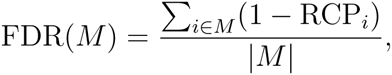

where |*M*| denotes the number of rejected hypotheses (Newton *et al.*, 2004, Müller *et al.*, 2004, Wen, 2016). Therefore, at a pre-defined FDR level *α*, the Bayesian FDR control procedure simply ranks all candidate loci according to increasing values of (1 − RCP_*i*_) and rejects the null hypotheses for the largest set *M*, where

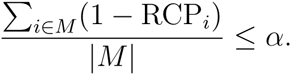

#### 2.4.3 Estimating Expected Value of Colocalized Association Signals

One of the useful utilities of the proposed hierarchical model and enrichment analysis is to estimate the expected number of colocalized association signals prior to delving into individual loci. The estimation is based on the proposed prior model and is analogous to the power calculation in, e.g., genetic association studies. We denote the marginal probabilities *p*_*γ*_ := Pr(*γ*_*i*_ = 1) and *p*_*d*_ := Pr(*d*_*i*_ = 1). Recall that 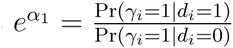, and it follows from simple algebra that

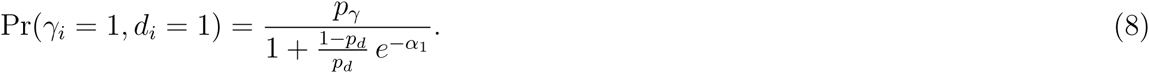

Note that the quantity

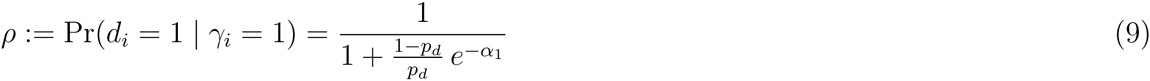

represents the the fraction of causal GWAS hits overlapping causal molecular QTNs.

The interplay of *p*_*d*_, *p*_*γ*_ and *α*_1_ with respect to *ρ* can be intuitively understood in some extreme scenarios. For example, if the vast majority of the genome is annotated as molecular QTNs, i.e., if *p*_*d*_ → 1, then *ρ* → 1 and Pr(*γ*_*i*_ = 1, *d*_*i*_ = 1) → *p*_*γ*_. This is because if every SNP in the genome is likely a molecular QTN, then every causal GWAS SNP is also likely a molecular QTN. More generally, the colocalization probability is impacted by the level of enrichment of molecular QTNs in the GWAS hits. Specifically, if *α*_1_ → ∞, ρ → 1 and Pr(*γ*_*i*_ = 1, *d*_*i*_ = 1) → *p*_*γ*_, i.e., all GWAS hits are expected to be molecular QTNs. Alternatively, if *α*_1_ = 0, it follows that *ρ* = *p*_*d*_ and Pr(*γ*_*i*_ = 1, *d*_*i*_ = 1) = *p*_*γ*_ *p*_*d*_, i.e., the two types of associations are mutually independent. Moreover, if molecular QTLs are depleted in the GWAS hits, i.e., *α*_1_ *<* 0, *ρ* is expected to be < *p*_*d*_.

To estimate the expected number of colocalized association signals, we simply compute

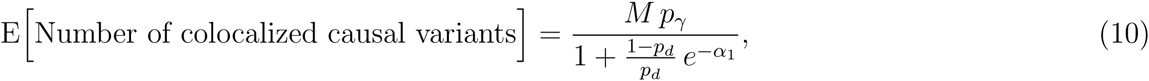

where *M* represents the total number of interrogated genetic variants.

## 3 Results

### 3.1 Simulation Study

First, we perform simulation studies to benchmark the performances of the proposed enrichment and colocalization analysis approaches.

We design the simulation scheme to generate realistic single SNP association *z*-statistics that are similar to the observed GWAS results. Specifically, we select real genotypes of 2.7 million overlapping SNPs used by both Wood *et al.* (2014) and the GTEx project from the European samples from the 1000 Genomes Project. For each SNP, we obtain its binary eQTL annotation by drawing from the posterior distribution of GTEx whole blood *cis*-eQTLs the GTEx. This particular posterior distribution is obtained by performing multi-SNP fine-mapping of the GTEx whole blood data via the adaptive DAP algorithm (Wen *et al.*, 2016). In total, we roughly annotate *∼* 6, 000 SNPs per simulation. We then simulate the association status of each SNP *i* (*γ*_*i*_) by drawing from a Bernoulli distribution whose success rate is determined by the logistic model (2) with pre-determined *α*_0_ and *α*_1_ values. Subsequently, a quantitative trait is simulated using a standard multiple linear regression model for which the residual error variance is set to 1 and the effect size of each causal SNP is drawn from a N(0, *ϕ*^2^) distribution. Finally, we compute the single SNP association *z*-statistic for each SNP as the input for both the enrichment and the colocalization analyses. Although the sample size in the 1000 Genomes Project European panel is limited, we are able to adjust the values of *α*_0_ (which determines the prevalence of the causal associations) and *ϕ* (which determines the signal-to-noise ratio of the genetic effects) to roughly match the *z*-value distributions from the available large-scale GWAS meta-analysis. In particular, we estimate *α*_0_ and *ϕ* by analyzing the height data reported in Wood *et al.* (2014), and we set *α*_0_ = −8.4 and *ϕ* = 0.4. Consequently, the distributions of the simulated *z*-statistics closely resemble the actual observed GWAS height data (Supplementary Figure S1). We vary the value of *α*_1_ across simulations for different levels of enrichment.

#### 3.1.1 Evaluation of Enrichment Analysis

We examine the performance of the proposed inference procedure in estimating the enrichment parameter *α*_1_. In particular, we vary the true *α*_1_ value in the range of 0.0 to 5.0 in the simulations. For each *α*_1_ value, we simulate 100 data sets and estimate *α*_1_ for each simulated data set using the proposed multiple imputation approach.

For comparison, we also estimate *α*_1_ using two additional approaches with added information. The first approach represents a best case scenario in which the true association indicators of each SNP in GWAS and eQTL mapping, i.e., *γ*_*i*_ and *d*_*i*_, are assumed to be observed. In this case, *α*_1_ is trivially estimated using a 2 × 2 contingency table. The second approach assumes that the association indicator of GWAS, *γ*_*i*_, is unobserved but that the true eQTL annotation for each SNP, *d*_*i*_, is known, which presents a type of integrative analysis considered in our previous work (Wen *et al.*, 2016). In this scenario, we apply the EM-DAP1 algorithm implemented in the software package TORUS (Wen, 2016) to estimate *α*_1_. Note that both of these approaches require additional information that is practically unattainable. Nevertheless, the results from these analyses highlight the intrinsic difficulty of the task and the theoretical ceiling of any realistic computational approach.

The results obtained using various estimation approaches are summarized in Figure 2. Importantly, note that when the enrichment level is low, the accurate estimation of *α*_1_ is difficult even in the best case scenario: the point estimates show large variance even when the true values of *γ*_*i*_ and *d*_*i*_ are known. In comparison, we observe that the estimates are significantly stabilized by applying the proposed adaptive shrinkage. As *α*_1_ increases to relatively large values (*>* 3.0), the effects of shrinkage gradually diminish for all approaches: in the case that the true QTL annotation is known, the estimates become practically unbiased, although for the multiple imputation procedure, the resulting estimates still notably biased toward 0. Nonetheless, we note that the degree of bias has minimal impact on the subsequent colocalization analysis (Figure 3).

**Figure 3:**
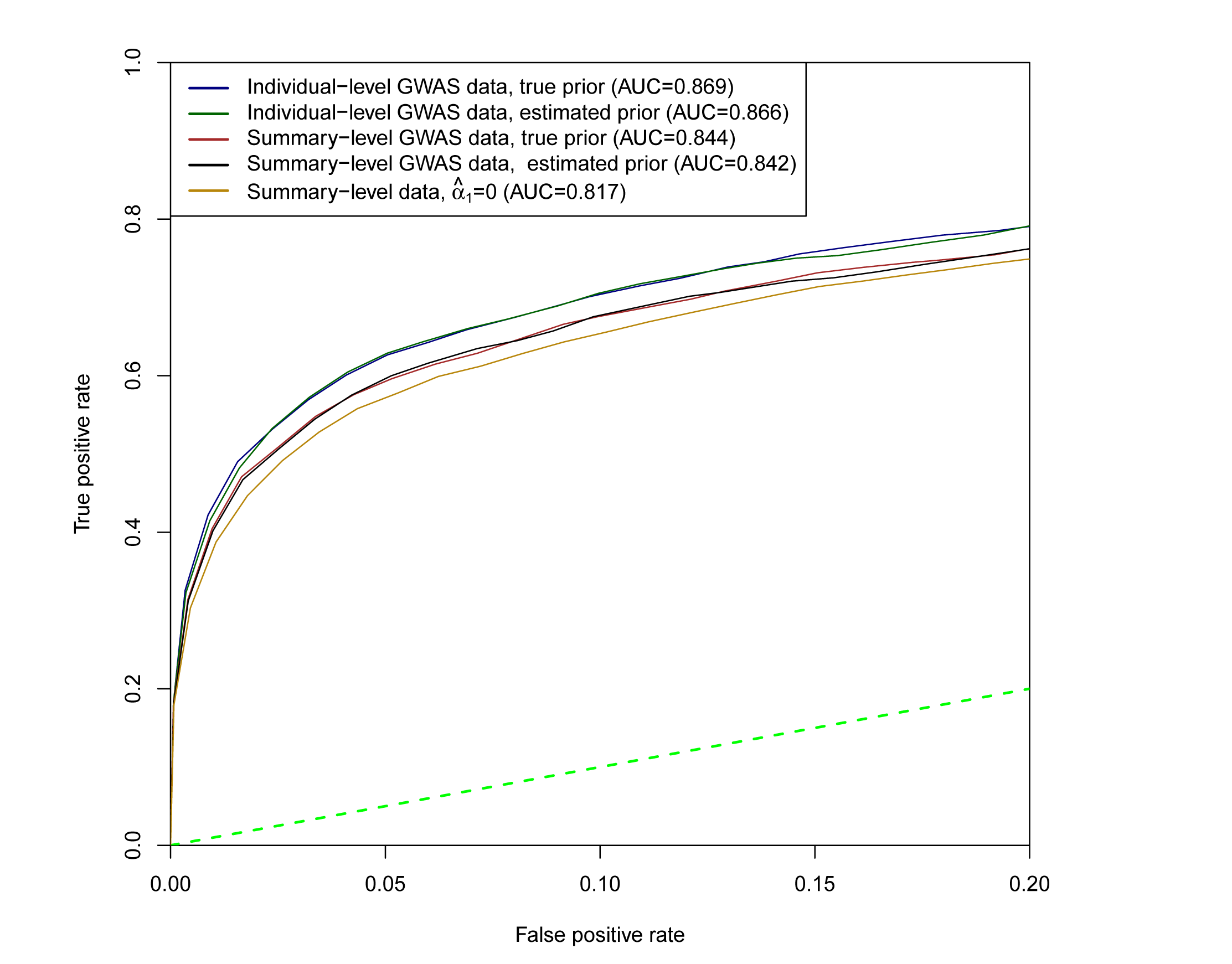
ROC curves for various colocalization analysis schemes in simulation studies. ROC curves evaluate the ranking of the LD blocks that potentially harbor colocalized association signals. All schemes perform decently in the simulations. Notably, the inaccuracy of the estimated enrichment parameters from the proposed multiple imputation procedure does not appear to have a significant impact on the overall performance of the colocalization analysis. However, the difference becomes highly visible for the case where *α*_1_ is set to 0. In addition, multi-SNP analysis in GWAS also improves the performance of the colocalization analysis.

**Figure 2:**
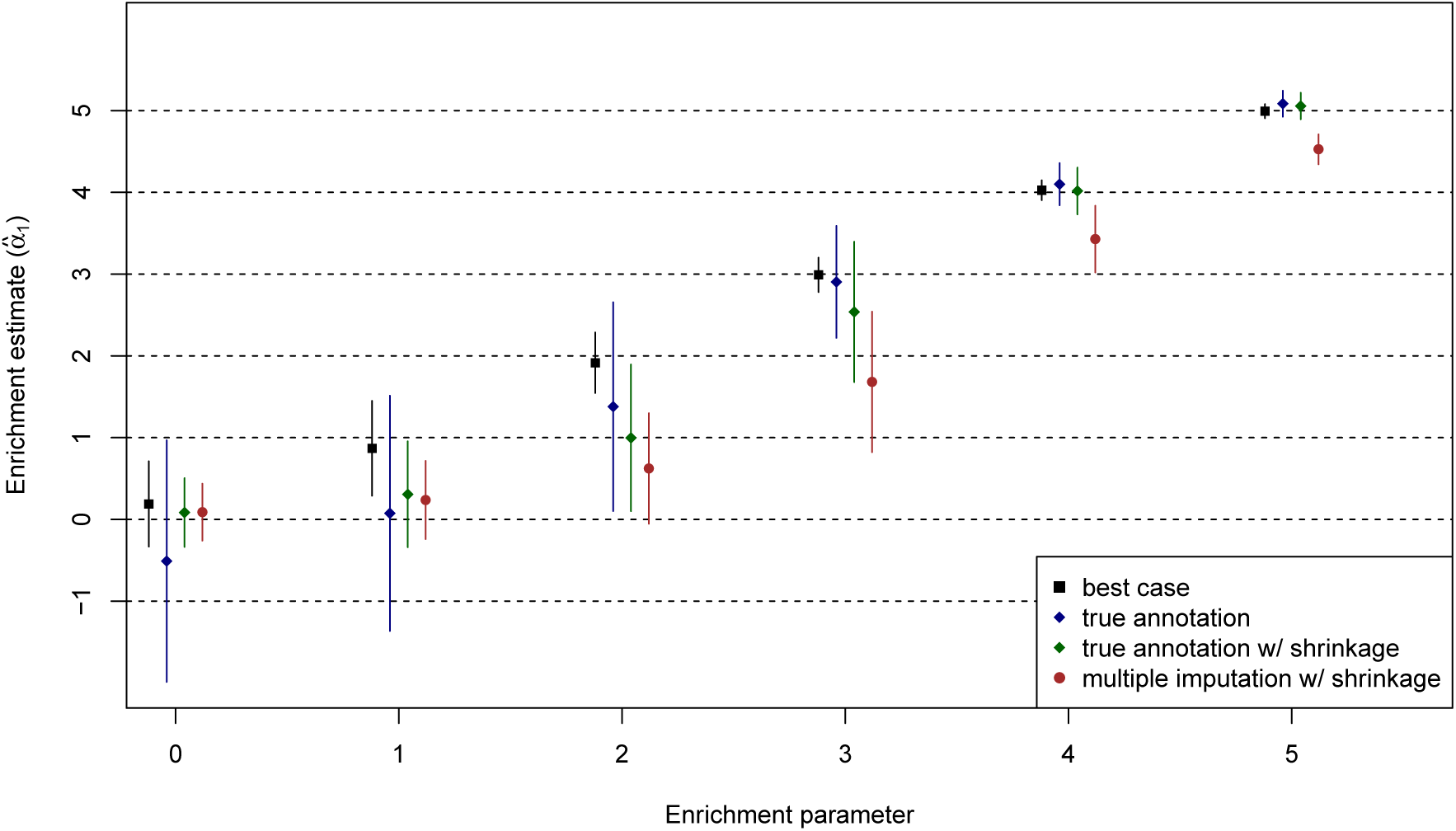
Enrichment parameter estimates in simulation studies. The proposed multiple imputation approach is compared to three methods utilizing added information that is unattainable in practice. The “best case” uses the true association status for both complex traits and molecular QTLs, whereas the “true annotation” utilizes the true association status from molecular QTLs only. This figure highlights the difficulty in estimating 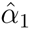 even when additional information is available. It shows the necessity of applying shrinkage to stabilize the point estimates in our simulation setting.

Because the overall accuracy of the estimate (rather than bias or variance individually) is more relevant to our downstream fine-mapping and colocalization analyses, we further investigate the precision of the estimate 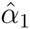 by computing the root-mean-square error (RMSE) for all methods. To this end, we also include two additional *ad hoc* imputation strategies for enrichment estimation. The first strategy applies “mean imputation”, i.e., for each SNP, we regard the marginal PIP of each SNP (which is also the posterior mean of the corresponding *d*_*i*_ value) as a continuous annotation. The second strategy, known as “best SNP imputation”, annotates the best associated *cis* candidate SNP of each eGene (i.e., the gene harboring at least one causal eQTL) as the causal eQTL. For all compared methods, we compute the RMSE based on the true *α*_1_ and estimated 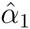 values across 600 simulated data sets. We find that the proposed multiple imputation approach outperforms both of the alternative imputation strategies in terms of the accuracy of the point estimates, and its precision is close to the case in which the true eQTL annotation is known (Table 1). It is highly expected that differences in performance will be observed between the multiple imputation and the best SNP imputation because the latter case ignores the uncertainty due to LD and the potential multiple independent eQTLs within a gene. We observe that the mean imputation approach consistently (and occasionally severely) overestimates *α*_1_, particularly for large *α*_1_ values, which becomes a serious concern for inflating type I errors in the downstream hypothesis testing of colocalization. In addition, the inability to apply the adaptive shrinkage (which is specifically designed for binary annotations) also contributes to the considerably larger RMSE result. Note that the use of mean imputation in our scenario is different than the case of mean genotype imputation commonly applied in GWAS. This is because in GWAS, the mean genotype imputation is typically highly accurate and there is also generally a stringent threshold for filtering out inaccurate imputation for downstream association analysis, whereas in our case, the PIPs are much less accurate representations for the true eQTL association status, particularly for true eQTNs.

**Table 1:**
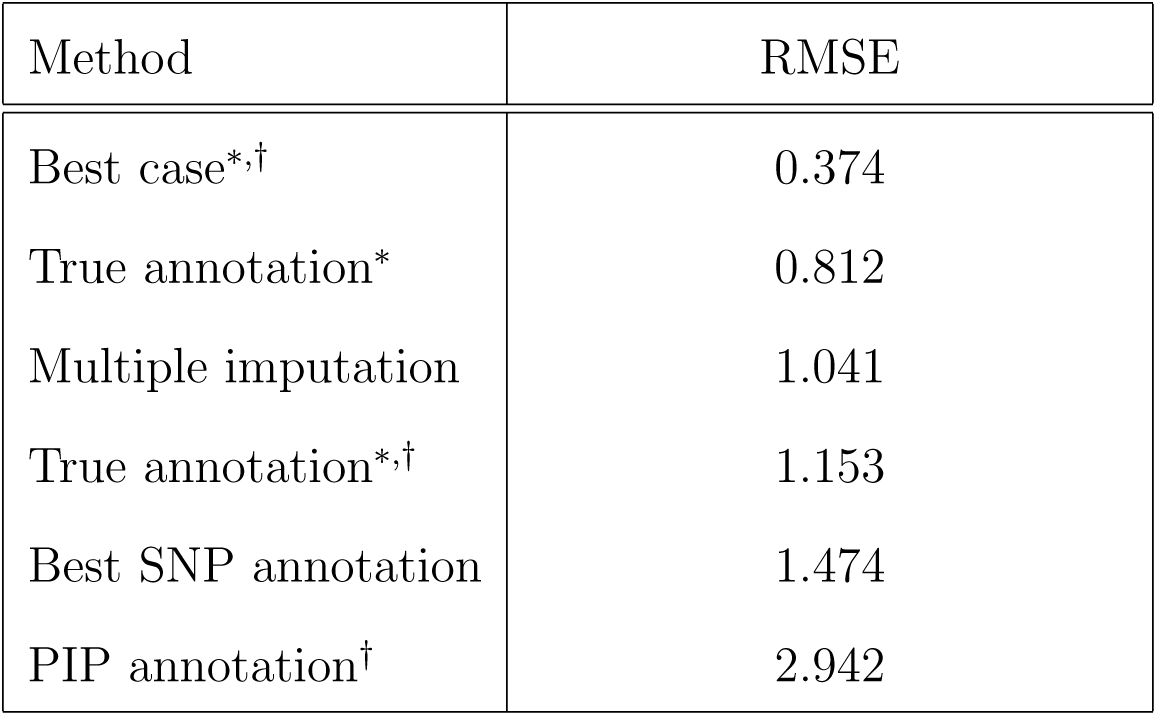
Evaluation of the accuracy of various enrichment estimation approaches. Using the simulated data sets, we compute the root-mean-square errors (RMSEs) to measure the precision of the point estimates obtained by different approaches. The methods denoted by ^*^ use added information that is unattainable in practice. The methods denoted by † do not apply shrinkage to the enrichment estimate. The proposed multiple imputation approach yields the best accuracy among approaches that are practically applicable.

| Method | RMSE |
| --- | --- |
| Best case<sup>*,†</sup> | 0.374 |
| True annotation<sup>*</sup> | 0.812 |
| Multiple imputation | 1.041 |
| True annotation<sup>*,†</sup> | 1.153 |
| Best SNP annotation | 1.474 |
| PIP annotation<sup>†</sup> | 2.942 |

In addition, we examine the statistical performance of the proposed approach for testing the null hypothesis

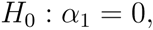

by inspecting the corresponding estimate of the 95% confidence interval from each simulated data set. Our results indicate that the testing results based on the proposed multiple imputation approach properly control type I error at the 5% level with the actual type error rate = 0.01. Although it achieves nearly perfect power as the true *α*_1_ *≥* 4, it only displays modest power (53%) for *α*_1_ = 3 and little power for smaller *α*_1_ values. Furthermore, despite the point estimates being downward biased, we observe that the proposed multiple imputation procedure provides excellent 95% interval estimates in the range of the *α*_1_ values examined experimentally: the coverage probability reaches 94.8%.

Finally, the benchmarked computational time indicates that the proposed multiple imputation approach is highly efficient. We take advantage of the fact that the multiple imputation scheme is parallelizable and analyze each simulated data set on 8 simultaneous threads. Consequently, each enrichment analysis only takes approximately 4 to 5 minutes of real computing time.

#### 3.1.2 Evaluation of Colocalization Analysis

To evaluate the performance of the colocalization analysis, we focus on the simulation setting of *α*_1_ = 4, which is close to our enrichment estimate of blood eQTLs in HDL GWAS hits. For each simulated data set, we perform the proposed colocalization analysis using two different finemapping strategies. The first strategy utilizes the individual-level genotype data from GWAS and obtains the GWAS PIPs by multi-SNP fine-mapping using the adaptive DAP algorithm. The second strategy assumes at most one causal GWAS hit within each LD block and computes the PIPs using the DAP-1 algorithm based only on the single-SNP association *z*-statistics. To evaluate the impact of (imperfect) enrichment parameter estimate, we separately use the true and estimated (*α*_0_, *α*_1_) values (by multiple imputations) to construct the SNP-level prior (5) for fine-mapping when applying each strategy. For comparison, we also perform the colocalization analysis of the simulated data assuming independence of molecular eQTLs and GWAS hits (i.e., set *α*_1_ = 0 in prior model (2)), which resembles *eCAVIAR* as shown previously. In all cases, we compute the RCPs for all the pre-defined LD blocks in each simulated dataset.

First, we construct receiver operating characteristic (ROC) curves to simultaneously evaluate the sensitivity and specificity of various colocalization analysis approaches. Specifically, we classify an LD block harboring a colocalized signal if the corresponding RCP is greater than a predefined threshold. We vary the threshold from 1 to 0 to construct the ROC curve for each analysis scheme. The results are presented in Figure 3, which highlights the performance of each examined approach as the corresponding false positive rates (FPR) *≤* 0.20. In summary, we find that all approaches yield decent results in identifying true colocalized signals while controlling for false positives (i.e., they are all well above the 45 degree diagonal line). In particular, we note that i) the ability to identify multiple independent GWAS hits within an LD block (i.e., in the adaptive DAP algorithm) improves the performance of colocalization analysis; ii) the downward bias in the enrichment parameter estimates by the proposed multiple imputation approach has very little impact on the colocalization analysis *at any given FPR threshold*; and iii) neglecting the enrichment analysis yields inferior colocalization results.

Note that the ROC curves rely only on the ranking of the corresponding RCPs and are invariant under the rank preserving transformations. To investigate the statistical properties of the RCPs reported by various analysis schemes as a probability measure, we further examine the Bayesian false discovery rate (FDR) control of colocalization analysis based on RCPs in the hypothesis testing setting described in the Method section. Figure 4 shows the comparison of estimated FDRs and the realized FDRs for all analysis schemes in the simulations. All approaches (conservatively) control the desired FDR levels; however, the scheme assuming *α*_1_ = 0 is extremely conservative, where the realized FDRs are nearly 0 and the power is significantly lower than all the other competing schemes. We therefore conclude that the accurate enrichment estimation has a critical impact on the quantification of the colocalized signals. In general, we find that the power to detect colocalized association signals is low across different schemes, i.e., *<* 40% at the 20% FDR level (Figure 4). Because our simulated data closely mimic the reality of the currently available GWAS and eQTL data, we attribute the lack of power reflected by these simulations to the limitations of the currently available genetic association data. (This point will be further demonstrated by the power calculation in the real data applications.)

**Figure 4:**
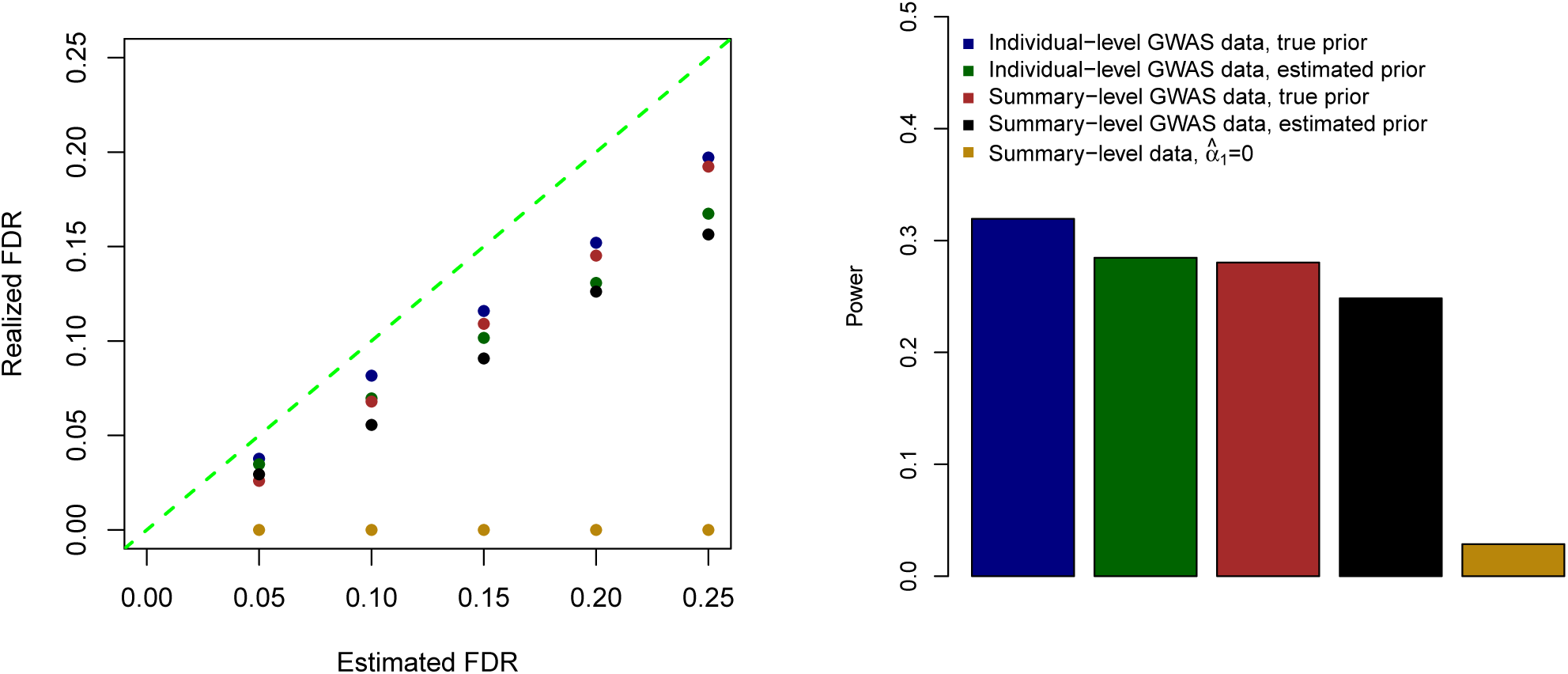
Evaluation of type I error rate and power for various colocalization analysis schemes in simulation studies. This exercise helps to evaluate the calibration of the reported RCPs from various analysis schemes. Better calibrated RCPs result in less conservative control of the type I errors and improved power. Our results indicate that the RCPs are better calibrated for more accurate enrichment estimates and/or the use of multi-SNP analysis in GWAS.

Taken together, we conclude that the estimation of the enrichment parameters embedded in the prior model (2) impacts both the ranking and calibration of locus-level posterior probabilities for colocalization. According to the ROC curves, the impact on the ranking can be relatively insignificant with respect to non-trivial deviation from the truth for the enrichment parameter. However, the calibration of the colocalization probabilities is considerably more sensitive to such deviation, as evidenced by the power and the realized FDRs in the hypothesis testing of colocalization.

### 3.2 Integrative Analysis of Blood eQTL and Lipid GWAS Data

To demonstrate the proposed computational approach in a practical setting, we perform the integrative analysis of the eQTL data from the GTEx project (Ardlie *et al.*, 2015) and the blood lipid data originally reported in Teslovich *et al.* (2010). The blood lipid data consist of meta-analysis results of four quantitative traits, including low-density lipoprotein (LDL) cholesterol, high-density lipoprotein (HDL) cholesterol, triglycerides (TG) and total cholesterol (TC), with an aggregated sample size of *∼* 100,000. We obtain the version of single-SNP association *z*-statistics for the four traits re-analyzed by Pickrell (2014), where additional *z*-statistics for untyped SNPs are imputed according to the 1000 Genomes Project phase I panel. In total, the complete data set contains *z*-scores of *∼* 6.1 million SNPs per trait. For most of our analysis, we focus on the *cis*-eQTL data from the whole blood in the recent release (version 6) of the GTEx project. The selection of the whole blood is informed by the consensus of multiple independent enrichment analysis approaches (GTEx consortium, manuscript in prep.) to determine the relevant tissues for the blood lipid traits. In addition to biological relevance, we suspect that one of the driving factors is that the whole blood is one of the GTEx tissues with the largest sample size (338) in the current release of the data; it therefore has better power to detect *cis*-eQTLs with small to modest effects. The SNPs that are not directly genotyped are also imputed according to the same 1000 Genomes panel by the GTEx consortium. We perform the Bayesian multi-SNP fine-mapping analysis for the GTEx whole blood data using the adaptive DAP algorithm and generate the joint posterior distribution Pr(***d*** | ***Y***_*qtl*_, ***G***_*qtl*_) while controlling for the SNP distance to the transcription start site (TSS) of the corresponding target gene. As shown in *Wen et al.* (2016), Wen (2016), this approach significantly improves the eQTL discovery.

#### 3.2.1 Expectation Calculation for Colocalization Analysis

Before conducting the proposed integrative analysis, we first compute the expected fraction of the GWAS hits of blood lipid traits that overlap blood *cis*-eQTLs using the approach described in the Method section. The calculation only requires an approximate estimate of the genome-wide prevalence of causal eQTLs. Here, we show two different approaches for obtaining the estimate.

The first approach utilizes the pre-computed posterior distribution of *cis*-eQTLs and calculates the expected fraction of eQTNs from the posterior distribution by

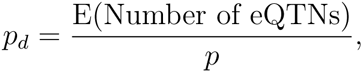

where the expected number of eQTNs can be conveniently obtained by summing up PIPs for all gene-SNP pairs. For the GTEx whole blood data, we calculate the posterior expected number of eQTNs as 8945.9, and hence, *p*_*d*_ ≈ 1.47 × 10^−3^.

Alternatively, we use a conservative *ad hoc* approach to estimate *p*_*d*_ without a Bayesian analysis of the *cis*-eQTLs. In particular, we note that the GTEx portal reports 6,784 eGenes (i.e., genes harboring *cis*-eQTLs) discovered in the whole blood samples at the 5% FDR level. Assuming that each eGene contains exactly one causal variant, we then estimate *p*_*d*_ ≈ 6,784/6.1 × 10^6^ = 1.11 × 10^−3^. Compared to the previous approach, which is more statistically rigorous, this estimate ignores potential multiple independent eQTNs within an eGene and the uncertainty embedded in the process of eGene discovery (e.g., a non-eGene could be mis-classified and indeed harbor eQTNs). Nevertheless, the two estimates have the same order of magnitude: roughly, we observe a causal *cis*-eQTL in 1 out of 1,000 SNPs.

We then calculate the expected fraction of GWAS hits overlapping causal eQTLs as a function of enrichment parameter *α*_1_ using the formula (9) for both estimates of *p*_*d*_. The result (shown in Figure 5) indicates that the expected fraction of overlapped signals is largely determined by the level of enrichment. With the current level of eQTL discovery (reflected by *p*_*d*_), we do *not* expect a large fraction of the GWAS hits to overlap with the annotated *cis*-eQTLs unless the enrichment level is extremely high. For example, even at *α*_1_ ∼ 5, which corresponds to a fold-change at ∼ 150, the expected fraction of colocalized GWAS signals is still less than 20% – in the case of the genetic variants associated with HDL, the expected number of colocalized signals is ∼ 10.

**Figure 5:**
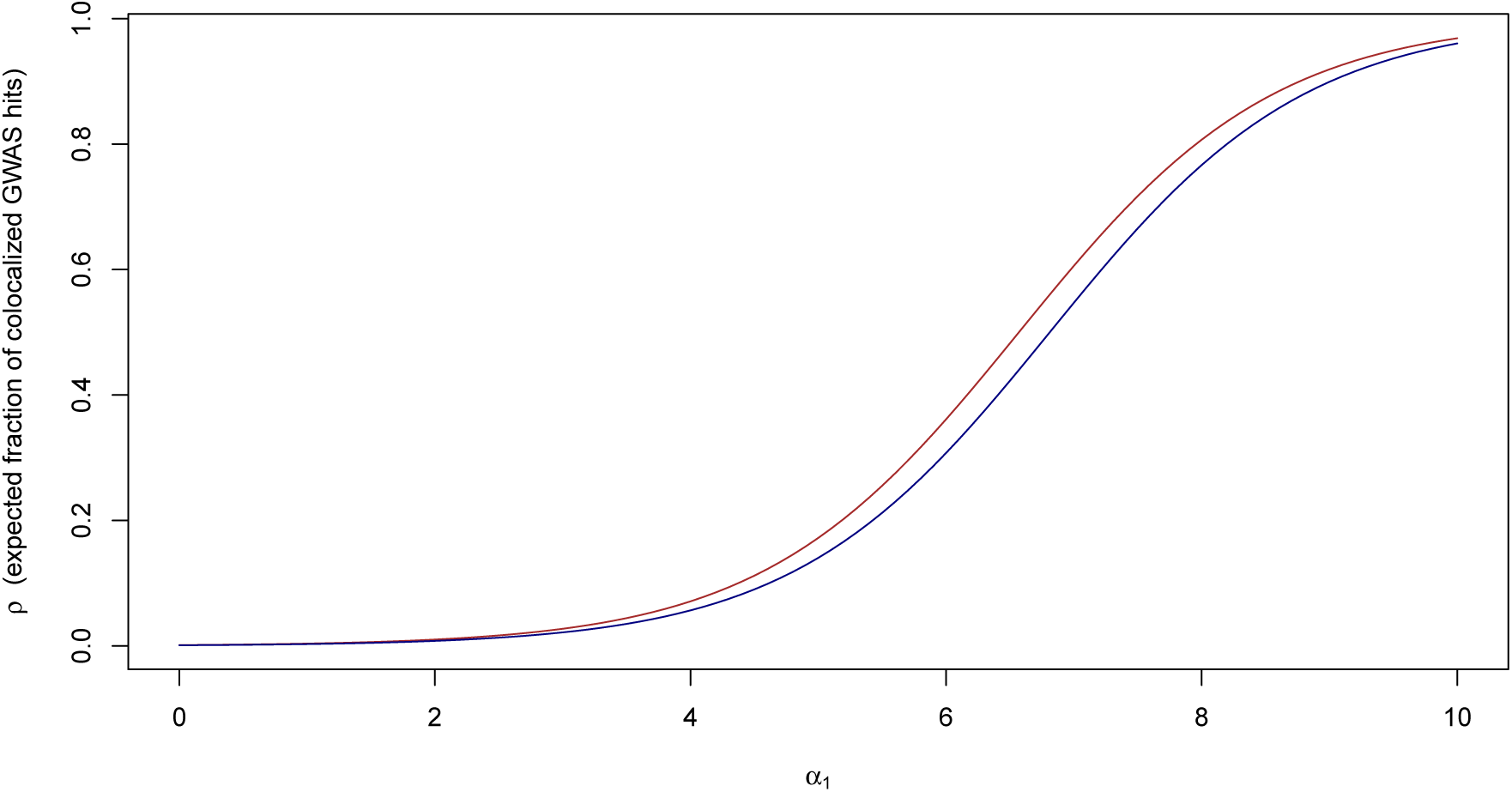
Expected fraction of colocalized GWAS hits in GTEx whole blood *cis*-eQTLs. The red and green curves are computed using the *p*_*d*_ estimates from a model-based and an *ad hoc* approach, respectively. Qualitatively, the two curves are similar. The expected fraction of GWAS hits overlapping *cis*-eQTLs is largely determined by the enrichment parameter *α*_1_: if *α*_1_ *→* 0, we should expect few colocalization signals, whereas if *α*_1_ is large, a large proportion of GWAS hits are expected to overlap with eQTLs.

#### 3.2.2 Enrichment Analysis

Next, we apply the proposed multiple imputation procedure to estimate the enrichment level of whole blood *cis*-eQTLs in the GWAS hits of the four lipid traits. As in the analysis of the simulated data sets, we apply the multiple imputation approach for each lipid trait by imputing 25 independent binary eQTL annotations from the joint posterior distribution of the blood *cis*eQTL data.

We show the enrichment estimates of blood *cis*-eQTLs in lipid traits and their corresponding 95% confidence intervals in Figure 6. We find that the blood eQTLs are most enriched in the GWAS hits of HDL with point estimate 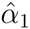 = 4.98, followed by TC (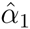 = 3.73), LDL (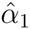 = 2.95) and finally TG (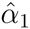 = 0.38). The behavior of the proposed enrichment analysis method is very consistent with what we observed in the simulation studies, i.e., imperfect power for enrichment estimates ≤ 4 as we note that the corresponding 95% confidence intervals cross 0. We further inspect the individual enrichment estimate from each imputed eQTL annotation for each trait (Supplementary Figure S2). We find that the enrichment estimates for HDL and TG are quite consistent across all imputed annotation data sets, whereas the estimates for HDL and TC show relatively large variations across imputed annotations. Nevertheless, we observe that all point estimates across all imputed annotations for all 4 traits are positive.

**Figure 6:**
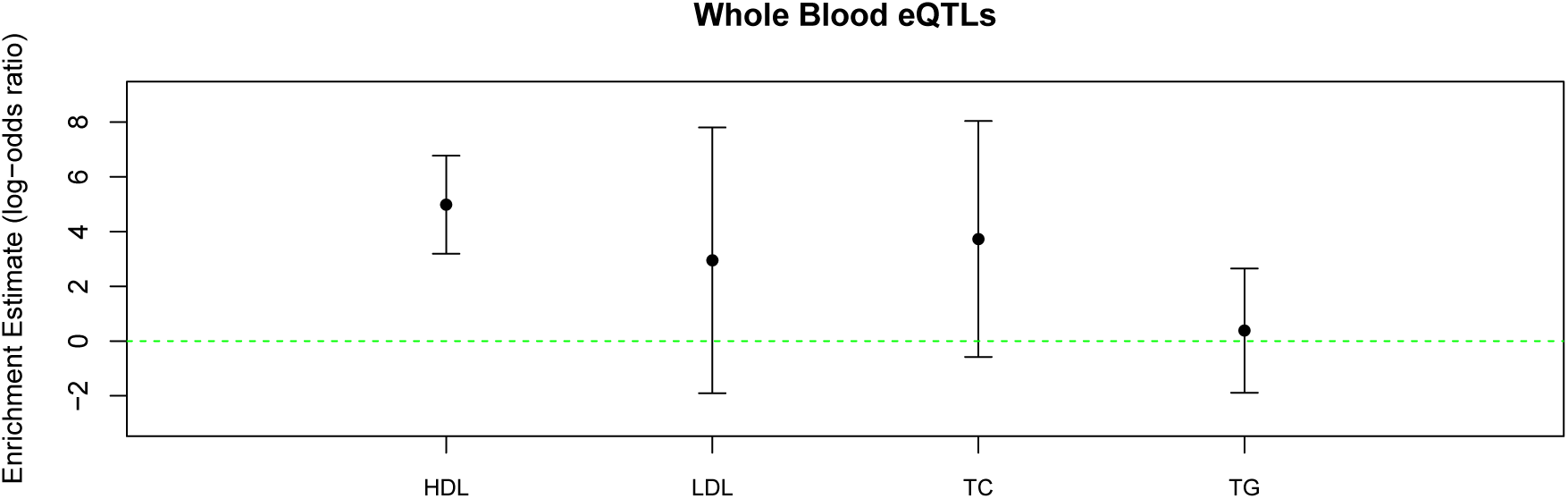
The enrichment estimate of GTEx whole blood *cis*-eQTLs in the GWAS hits of four blood lipid traits. For each trait, the point estimate and the corresponding 95% confidence interval are plotted.

We then plug in the enrichment estimates and calculate the expected fraction of colocalized GWAS hits for each trait from the previously constructed power curves. In summary, we expect that 18%, 3%, 6% and 0.2% of GWAS hits in HDL, LDL, TC and TG overlap with causal *cis*-eQTLs in whole blood. Although the true fractions of overlaps may have large variations due to the uncertainty embedded in the enrichment estimates (as indicated by their large CIs), these estimated expected fractions should reflect the relative difficulty in finding colocalized signals in each lipid trait.

#### 3.2.3 Colocalization Analysis

Given the enrichment estimates, we proceed to perform the colocalization analysis. Because the blood lipid data are provided in the form of summary statistics, we adopt the “signal-centric” approach that focuses on colocalizing the association signals of lipid traits identified in the single-SNP association analysis. Specifically, we apply the Bayesian FDR control procedure (Wen, 2016) implemented in TORUS to identify the LD blocks (defined in Berisa and Pickrell (2016)) that harbor at least a single association signal at the 5% FDR level. Ultimately, we identify 72, 64, 78 and 52 genomic loci for HDL, LDL, TC and TG, respectively. Because Teslovich *et al.* (2010) controlled for the family-wise error rate (FWER) and used a stringent SNP-level genome-wide significance threshold (i.e., 5 × 10^−8^), their reported loci consist of a subset of ours.

Another practical issue arising in the eQTL analysis is that a single SNP can locate in the *cis* regions of multiple genes. Our solution to this problem is to compute an RCP for each LD block with respect to each gene that has at least one *cis* candidate SNP residing in the block. In particular, we construct the SNP prior (5) that is specific to each gene. Consequently, the resulting RCP of the target LD block is also gene specific. One obvious advantage of this strategy is that it naturally links the SNP-level association signal to a gene.

In summary, our analysis identifies 21 unique genomic region-gene pairs with RCP ≥ 0.50. We summarize the results in Table 2. In the hypothesis testing context, we reject 4 (RCP cutoff 0.902), 7 (RCP cutoff 0.832) and 16 (RCP cutoff 0.639) top-ranked RCP regions at the Bayesian FDR levels of 5%, 10% and 20%, respectively. Within an LD block, we regard an SNP as a potentially true colocalized signal if its SCP is ≥ 0.001.

**Table 2:** Identified genomic regions that potentially harbor colocalized association signals of whole blood *cis*-eQTLs and GWAS hits of blood lipid traits. A region is listed if its RCP is ≥ 0:5. We denote the region by * if it is also identified in Teslovich *et al*. (2010). The symbol † indicates that the same gene is linked to the same GWAS hit region in Teslovich *et al*. (2010).

| Trait | Region | Gene | RCP | # of SNPs | Lead SNP | Max SCP |
| --- | --- | --- | --- | --- | --- | --- |
| HDL | chr1:109817192-109818530* | *PSRC1* | 0.962 | 5 | rs629301 | 0.439 |
| HDL | chr2:85537312-85555478 | *AC093162.5* | 0.845 | 22 | rs10460586 | 0.340 |
|  |  | *TCF7L1* | 0.828 | 27 | rs10460586 | 0.208 |
|  |  | *ELMOD3* | 0.800 | 22 | rs3184780 | 0.099 |
| HDL | chr3:49971514-50146094 | *RBM6* | 0.712 | 22 | rs7613875 | 0.380 |
| HDL | chr6:34548206-34800435* | *C6orf106*<sup>†</sup> | 0.814 | 35 | rs6907508 | 0.623 |
| HDL | chr9:15303583-15304782* | *TTC39B*<sup>†</sup> | 0.627 | 2 | rs686030 | 0.580 |
| HDL | chr11:61557803-61623140* | *TMEM258* | 0.832 | 15 | rs102275 | 0.584 |
| HDL | chr11:122500846-122553139* | *UBASH3B*<sup>†</sup> | 0.639 | 54 | rs60494825 | 0.036 |
| HDL | chr12:109893156-110042348* | *MVK*<sup>†</sup> | 0.603 | 45 | rs7964021 | 0.051 |
| HDL | chr19:54796630-54799083* | *LILRA3*<sup>†</sup> | 0.990 | 3 | rs103294 | 0.979 |
| HDL | chr22:21917450-21980894* | *UBE2L3*<sup>†</sup> | 0.554 | 39 | rs181360 | 0.052 |
| LDL | chr1:109818306-109818530* | *PSRC1* | 0.901 | 2 | rs629301 | 0.879 |
| LDL | chr9:136141870-136155000* | *ABO*<sup>†</sup> | 0.582 | 5 | rs550057 | 0.430 |
| LDL | chr17:8107979-8161149 | *C17orf44* | 0.708 | 6 | rs8078338 | 0.637 |
| TC | chr1:109817590-109818530* | *PSRC1* | 0.942 | 4 | rs629301 | 0.858 |
| TC | chr9:136141870-136155000* | *ABO*<sup>†</sup> | 0.509 | 4 | rs635634 | 0.327 |
| TC | chr17:8107979-8161149 | *C17orf44* | 0.745 | 5 | rs8078338 | 0.671 |
| TC | chr19:49206108-49219459* | *NTN5* | 0.662 | 7 | rs492602 | 0.177 |
| TC | chr20:34124336-34160840* | *RPL36P4* | 0.745 | 15 | rs2277862 | 0.494 |

For a small proportion of the identified loci, we find that the colocalized signals can be effectively narrowed down to only a few SNPs. For example, SNP rs103294, a *cis*-eQTL for gene *LILRA3*, has an SCP value of 0.979, showing a strong SNP-level colocalization signal with the GWAS hit of HDL. However, the majority of the loci still carry many candidate SNPs due to common LD patterns present in the genetic data of both complex traits and molecular phenotypes. Figure A1 illustrates a colocalized association signal for HDL and the expression of *UBASH3B* in a 52 kb genomic region on chromosome 11. Our analysis identifies 54 SNPs with a joint RCP = 0.645; however, the strongest individual SCP is merely 0.036. Additionally, the SNP-level PIPs for GWAS and *cis*-eQTL associations also exhibit a similar pattern: although there is not a single SNP taking high PIP values, the cumulative PIPs of the region are close to 1 for both GWAS and *cis*-eQTLs. We further compute the pair-wise LD of the 54 member SNPs based on the genotype data from the GTEx samples and confirm that all SNPs are indeed highly correlated.

For the significant loci reported by Teslovich *et al.* (2010) (labeled by ^*^ in Table 2), we compare the genes suggested by our analysis and the reported genes therein. For 8 out of 14 cases, the implicated genes are consistent (labeled by ^†^ in Table 2). Among the other 6 inconsistent cases, 3 involve the genomic region anchored by SNP rs629301, for which our analysis links to *PSRC1* and Teslovich *et al.* (2010) links to *SORT1* by the more comprehensive molecular evidence presented in Musunuru *et al.* (2010). Our examination of the current GTEx analysis results (version 6) reveals that rs629301 shows little to no evidence of association with *SORT1* but very strong evidence of association with *PSRC1* in whole blood; however, in liver, rs629301 shows strong associations with both genes with evidence for *SORT1* being stronger (source: GTEx portal eQTL browser). In addition, the very SNP also shows strong association with *CELSR2* in liver. We repeat the colocalization analysis using the GTEx liver eQTL data. Not surprisingly, we find that the same genomic region presents the strongest colocalization signals with all 3 genes among all liver-expressed genes, with RCPs = 0.691, 0.684 and 0.675 for *SORT1*, *PSRC1* and *CELSR2*, respectively. The decrease of the RCP values is attributed to the lower eQTL enrichment estimate in liver (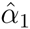 = 2.567 with 95% CI [−1.849, 6.984]), which exhibits a considerably larger degree of uncertainty than the whole blood estimate and is likely caused by the insufficient sample size in the current GTEx liver data (sample size of 97 compared to 338 for whole blood). Additionally, we find that the other 3 inconsistent cases can be similarly explained: the blood eQTLs for genes *RPL36P4*, *NTN5* and *TMEM258* all display different association patterns in different types of tissues.

Finally, we note that a single GWAS association can be colocalized to eQTL signals of multiple genes. For example, our analysis indicates that the likely causal HDL variant in the genomic region chr2:85537312-85555478 is possibly associated with the expression levels of 3 different genes (*AC093162.5, TCF7L1* and *ELMOD3*). The case of SNP rs629301 in liver discussed previously is also an example of this type. Although this phenomenon is relatively well known in studies of molecular phenotypes, it certainly makes elucidating the molecular mechanism of causal GWAS variants more challenging.

## 4 Discussion

In this paper, we have proposed a statistically rigorous and computationally efficient analytic framework for performing integrative analyses of GWAS and molecular QTL data and providing quantitative assessments of enrichment and colocalization of their association signals. One of the intrinsic challenges in genetic association analysis is that the resolution of identified association signals is always limited by LD. Consequently, it is generally impossible to pinpoint the causal variants based solely on genetic association analysis, and it imposes a formidable challenge for assessing enrichment and colocalization in the integrative analysis. To address this problem, we formulate a missing data problem and adopt a well-established statistical strategy, i.e., multiple imputation, to fully account for the uncertainty in identifying causal genetic variants for complex traits and molecular phenotypes due to LD. These efforts result in not only more accurate point estimates but also appropriate characterizations of uncertainties of our inference results in enrichment and colocalization analysis. Particularly, in the colocalization analysis, our theoretical demonstration and the real data example both clearly illustrate that individual SCPs can be unimpressive in high LD regions even if we are confident that the region does harbor a colocalized signal. In light of these findings, we propose and recommend reporting RCPs rather than placing emphasis on colocalization probabilities of individual SNPs.

Compared to the existing methods for colocalization analysis, the most important distinction of our proposed approach is the natural integration of the enrichment estimation. Throughout the paper, we have illustrated the importance of obtaining accurate enrichment estimates on the downstream quantitative evaluations of colocalization. Our main conclusion is that the accurate enrichment estimates based on currently available data may not have an overall large effect on altering the ranking of potential colocalization signals; however, it is critically important for the calibration of the corresponding colocalization probabilities and has a profound impact on the outcome of formal statistical testing procedures. Existing probabilistic model-based approaches typically make explicit assumptions on the enrichment levels of molecular QTLs in the causal GWAS hits (although they may not be presented in the form of enrichment parameters), as we have shown for the cases of *coloc* and *eCAVIAR*. We further hypothesize that all approaches, including empirical methodologies, make implicit assumptions on the enrichment parameters, which can be understood by the hypothetical example of two perfectly linked SNPs discussed in the Method section. For example, if a method determines that the association signals are colocalized in the hypothetical example (without enrichment estimation), it seemingly assumes that the enrichment level → ∞, which is a strong assumption. In summary, we have demonstrated that the enrichment parameter plays an important role in colocalization analysis, and we believe that the best strategy to deal with this parameter is to learn it from the observed data, as we have demonstrated throughout.

Importantly, our simulation and real data analyses apparently illustrate the limitations of currently available association data: we have shown that the confidence intervals of enrichment estimates are typically large and the expected fractions of colocalized GWAS signals are only modest, which are consistent with our observations from practice in the field. In particular, we note that most current molecular QTL (e.g., eQTL) studies are conducted with only modest sample sizes due to cost considerations. Although many of these studies successfully identified an abundance of trait-associated genomic loci with large effects, the power required to uncover molecular QTLs with small to modest effects is lacking. As many molecular QTL studies have started scaling up their sample sizes, we expect an elevated estimate of *p*_*d*_ in the future. Accordingly, based on Equation (9), we anticipate that a higher fraction of GWAS hits overlapping molecular QTLs can be revealed. Similarly, improving the power of GWAS should also help improve discoveries of colocalized signals, which is evident from Equation (10).

Our proposed statistical model and inference procedure are completely general for analyzing two sources of genetic association data. Note that it is statistically equivalent to treating GWAS data as annotations for eQTL mapping. Our choice of presenting eQTL as an annotation is simply motivated by better biological interpretation of the model and our enrichment analysis. It is trivial to prove that when individual-level data are available for both eQTL mapping and GWAS analysis, the choice of annotation should not alter the inference results under the proposed model. More generally, the proposed statistical framework is applicable for analyzing any pair of phenotypes to colocalize the association signals, as in applications demonstrated by Pickrell *et al.* (2016).

Note that caution should be exercised when attempting to interpret the biological relevance of the identified colocalization signals. In colocalizing an eQTL and a GWAS hit, a seemingly obvious implication is the relevance of the target gene of the eQTL in the disease process. However, as we demonstrated in the analysis of the blood lipid data, there are cases in which other important biological factors should be considered: for example, the relevance of the tissue where the eQTLs are derived from. Although it is generally possible to statistically evaluate the relevance of eQTLs from different tissues for a specific complex trait through enrichment analysis, the currently available GTEx data are not satisfactory for this purpose because of the significant variations in sample size across tissues. (We anticipate that this issue will likely be resolved by the end of the GTEx project, and we should re-visit the problem then.) A more elegant solution is to utilize eQTL annotations generated from joint multi-tissue eQTL mapping approaches (Flutre *et al.*, 2013, Li *et al.*, 2013), which enables simultaneous colocalization analysis across multiple tissues. Although conceptually straightforward, the difficulty in implementing a computationally efficient procedure incorporating multi-tissue eQTL data should not be underestimated. We will address this challenge in our future work.

In Teslovich *et al.* (2010), the authors went to great lengths to establish the biological, clinical and population relevance of genomic loci uncovered in the GWAS, in which integrative genetic association analysis is only a part of the overall process. Despite its own importance, we should acknowledge that integrative analysis of genetic association data is merely a starting point for uncovering the molecular pathway from genetic variants to complex traits.

## Acknowledgments

We thank the GTEx Consortium and the Global lipids Genetic Consortium for releasing valuable data in a timely fashion.

## 6 Resources

The URLs for data presented herein are as follows:

- Software pipeline and utilities for integrative analysis, http://github.com/xqwen/integrative/
- GTEx data, http://www.gtexportal.org/home/datasets
- Global Lipids Genetic Consortium results (2010), http://csg.sph.umich.edu//abecasis/public/lipids2010/

## Appendices

## Appendix A Multiple Imputation Procedure

In this section, we outline the multiple imputation procedure used for estimate *α*_1_.

We first analyze the GTEx whole blood *cis*-eQTL using the adaptive DAP algorithm, and obtain the joint distribution of Pr(***d*** | ***Y***_*qtl*_, ***G***_*qtl*_) for each gene. Subsequently, we create *m* = 25 imputed annotations by independent sampling from the corresponding posterior distribution of each gene. We then perform the enrichment estimate for each imputed data set using the EMDAP1 algorithm implemented in the software package TORUS. In the end, we obtain a point estimate of *α*_1_, namely 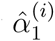, and its standard error, *σ*^(*i*)^ for each imputed data set *i*.

The multiple imputation procedure combines the estimates from individual imputed data sets in the following way. First, the overall point estimate 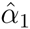 is simply given by

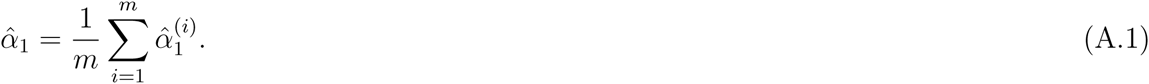

The variance of the point estimate is then computed by

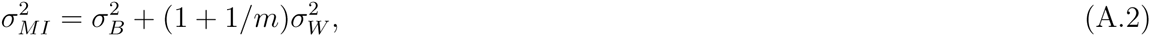

where

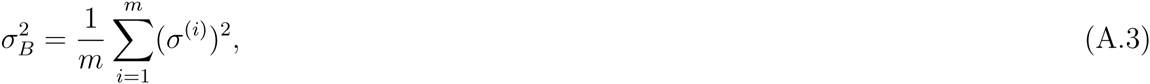

and

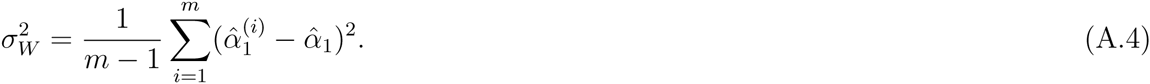

A more detailed reference on multiple imputation procedure can be found in Schafer (1997).

## Appendix B Adaptive Shrinkage in Enrichment Estimation

As discussed in the main text, the lack of strongly colocalized signals leads to unstable enrichment estimate. Here we propose a data-driven empirical approach to remedy this issue. The general statistical idea is to trade off the variance of the enrichment estimate against bias to achieve an overall accurate prior estimate for the downstream fine-mapping and colocalization analyses.

In the EM algorithm detailed in Wen *et al.* (2015) and Wen (2016), we showed that the M-step is equivalent to fitting a logistic regression model, which, in our context, regress the expected association status estimated in the E-step on the imputed eQTL annotation for each SNP. To apply the shrinkage on the estimate, we apply an *l*_2_ penalty with a constant shrinkage parameter *λ* in fitting the logistic regression for each M-step, which is equivalent to assuming a normal prior N(0, 1*/λ*) on *α*_1_.

We determine the shrinkage parameter in a data-adaptive way. Intuitively, a larger shrinkage is desired if the data indicates more severe sparsity of overlapping association signals. If both ***γ*** and ***d*** are observed, we can construct a 2 × 2 contingency table and *α*_1_ and its variance can be estimated in a standard way. In particular, the variance of the log-odds ratio estimate can be used as a measure for our purpose: e.g., the variance is small if all 4 cells have sufficient counts, whereas if the count of one cell is small, the overall variance of the log-odds ratio estimate is large. For each imputed data set, ***d*** is indeed observed but ***γ*** remains latent. We therefore estimate the cell count of the hypothetical 2 × 2 contingency table by computing the PIP of each SNP in GWAS using the DAP-1 algorithm and assuming *α*_1_ = 0. More specifically, let *η*_*i*_ denote the indicator if the SNP *i* is a molecular QTN in the imputed data set and let *p*_*i*_ denote the resulting PIP from the DAP-1 algorithm. Assuming there are total *p* candidate SNPs, we estimate the 4 cell counts by

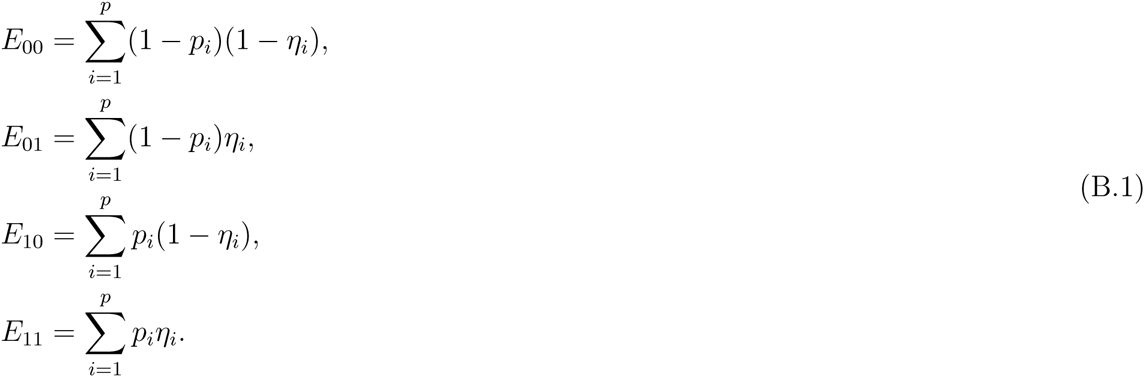

Finally, we set the shrinkage parameter *λ* by

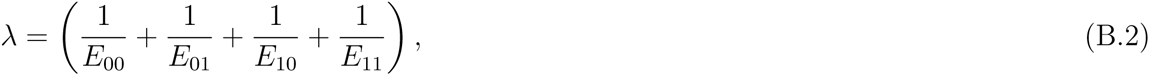

which is the variance of the log-odds ratio estimate if true cell counts are indeed observed (instead of estimated).

## Appendix C Derivation of SNP-level Colocalization Probability

The SNP-level colocalization probability can be computed by noting the relationship between the posterior probabilities Pr(*γ*_*i*_ = 1, *δ*_*i*_ = 1 | ***y***, ***G***, ***Y***_*qtl*_, ***G***_*qtl*_, 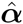) and Pr(*γ*_*i*_ = 1, *δ*_*i*_ = 1 | ***y***, ***G***, ***Y***_*qtl*_, ***G***_*qtl*_, 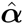) and their relationship to the marginal PIP in the GWAS data, i.e.,

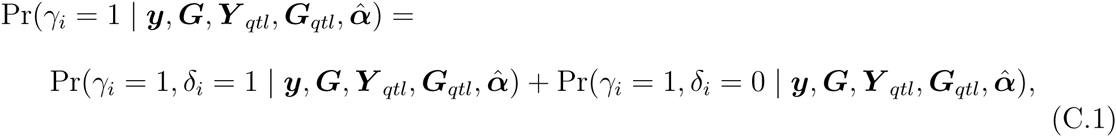

and

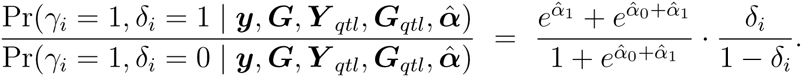

Solving this linear system results in the SNP-level colocalization probability expression (6).

## Appendix D Connection to *coloc* Model

Here we compare the statistical approach *coloc* to our proposed method. Specifically, we show that *coloc* can be viewed as a rough approximation and a special case to our general integrative analysis approach.

The method *coloc* requires pre-defining three SNP-level prior probabilities *p*_1_, *p*_2_ and *p*_12_. Using the notations of this paper, these three quantities can be formulated as

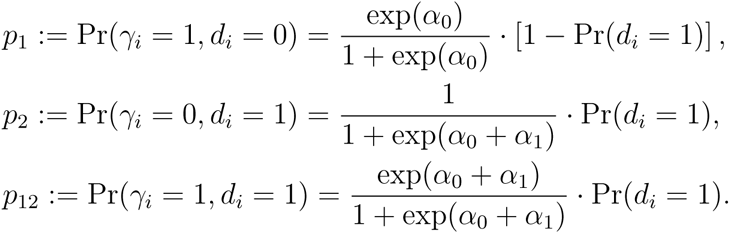

Additionally, the prior probability Pr(*γ*_*i*_ = 0, *d*_*i*_ = 0) = 1 − *p*_1_ − *p*_2_ − *p*_12_ can be trivially computed and represented by *p*_0_. In comparison, we explicitly estimate *α*_0_ and *α*_1_ from the GWAS data. Although we do not directly utilize the prior probability of a QTN, Pr(*d*_*i*_ = 1) (rather, the inferred posterior probability Pr(*d*_*i*_ = 1 | ***Y***_*qtl*_, ***G***_*qtl*_) is applied throughout our inference procedure), this very quantity can be straightforwardly estimated from the eQTL data, (***Y***_*qtl*_, ***G***_*qtl*_), using the EM-DAP1 algorithm (Wen *et al.*, 2016).

Similar to our RCP quantification, the *coloc* method considers the existence of a colocalized GWAS and molecular QTL signal within a LD block. Importantly the *coloc* model makes an explicit assumption that there is at most a single GWAS hit and/or a single QTN within the locus of interest, which enables highly efficient approximate computation for the RCP. Here we show that, given the simplifying assumption and the pre-specified priors for *p*_1_, *p*_2_ and *p*_12_, *coloc* yields identical result of RCP as the proposed method given *p*_1_, *p*_2_ and *p*_12_.

Suppose that there are *m* SNPs in the LD block of interest and let the binary *m*-vectors ***γ***_*l*_ and ***d***_*l*_ denote their association status with respect to the complex trait and the molecular phenotype, respectively. Assuming that SNP *l*_*i*_ is *the* colocalized association signal, it follows from our proposed model that

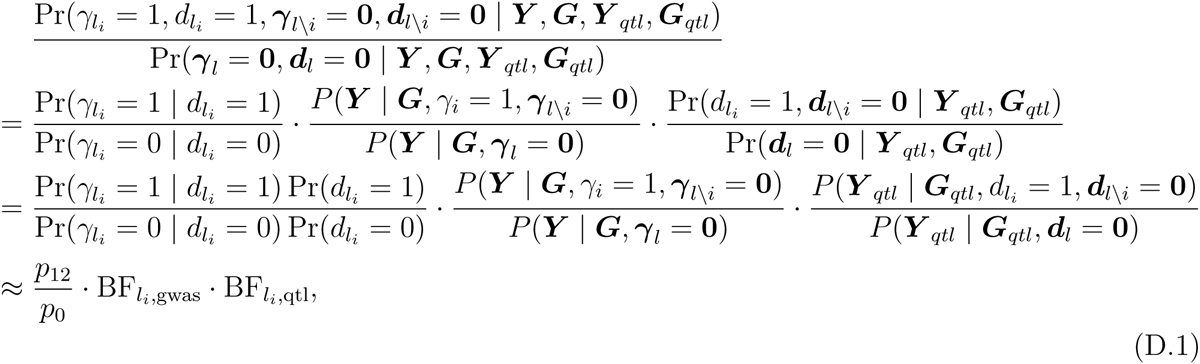

where the marginal likelihood ratios are approximated by the Bayes factors of single-SNP association models for the complex trait and molecular phenotype, respectively.

Under the constraint imposed by the simplifying assumption, each possible configuration of (***γ***_*l*_, ***d***_*l*_) can be enumerated, of which the corresponding posterior probability can be similarly computed as (D.1). For example,

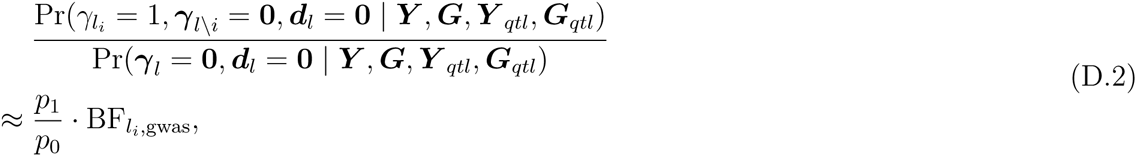

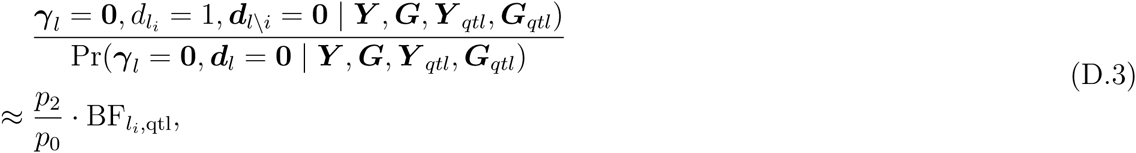

and,

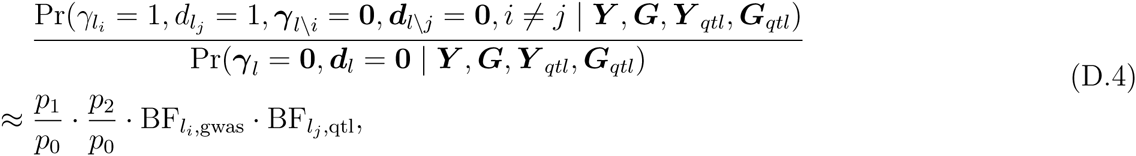

In the end, *coloc* computes the RCP for the locus of interest by

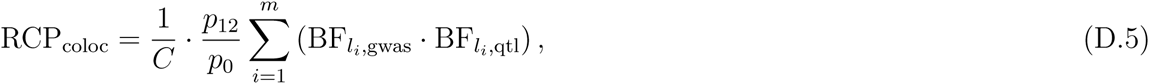

where *C* denotes the normalizing constant and is computed by

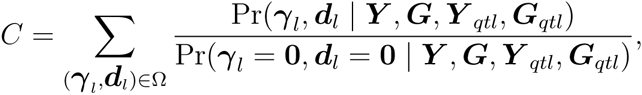

and the set Ω denotes the eligible (***γ**l, **d**_l_*) configurations under the constraint.

In summary, we have shown that the *coloc* approach can be derived as a special case from our proposed general framework with the following added assumptions,

1. the prior values are pre-defined through *p*_1_, *p*_2_ and *p*_12_
2. there is at most one causal GWAS variant in the region of interest
3. there is at most one causal molecular QTN in the region of interest

## Appendix E Multi-SNP Analysis with Adaptive DAP Algorithm using Summary-level Statistics

For practical and/or privacy considerations, many GWAS data are made available with only summary-level statistics, typically in the forms of single-SNP testing *p*-values or *z*-scores. In this section, we discuss the analytic strategy to perform proposed analysis using only summary-level statistics from GWAS. Many authors have demonstrated that the SNP-level PIP in GWAS, Pr(*γ*_*i*_ = 1 | ***y***, ***G***, ***Y***_*qtl*_, ***G***_*qtl*_, 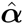), can be *approximated* from the summary-level statistics obtained from single-SNP association testing results (Kichaev *et al.*, 2014, Zhu and Stephens, 2016). Here we show their results extend to the application of the adaptive DAP algorithm.

In the case where individual-level genotype data are unavailable, the main computational difficulty in applying adaptive DAP algorithm lies in calculating the marginal likelihood, i.e., Bayes factor, for any given value of ***γ***. The evaluation of the Bayes factor or marginal likelihood using summary-level statistics has been systematically studied in the literature. In particular, we find the results by Chen *et al.* (2015) (Equation (3)) and Zhu and Stephens (2016) (Equation (2.10)) are suitable approximations that are accurate and easy to compute. With these results, it then becomes straightforward to apply the adaptive DAP algorithm to compute the PIPs for GWAS using the informative priors that incorporate molecular QTL annotaions.

## Figure Legends

**Figure A1:**
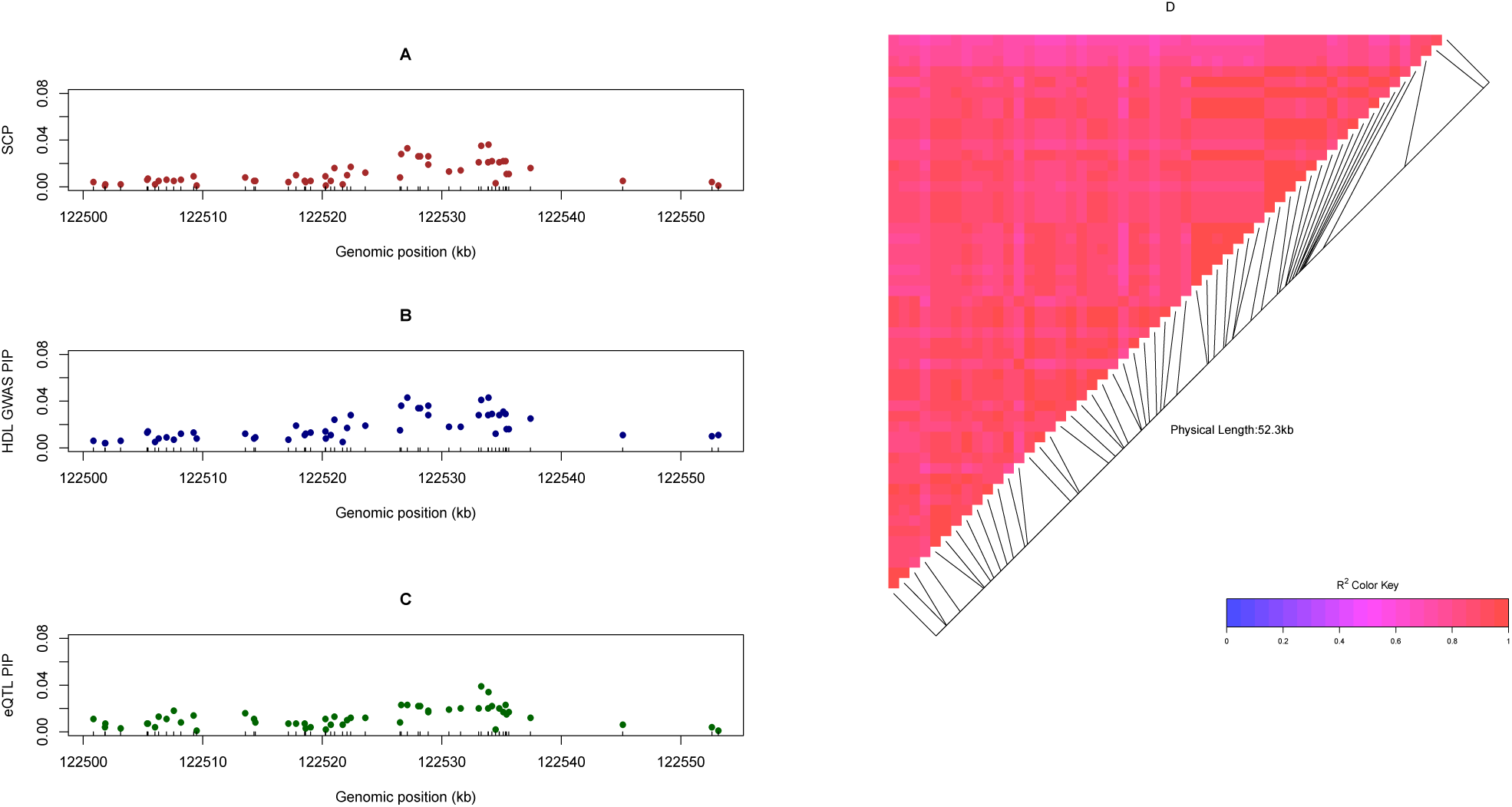
An example of an identified colocalization signal in a high LD region. The region, containing 54 candidate *cis*-eQTL SNPs for gene *UBASH3B*, harbors a GWAS hit for HDL. Panels A, B and C plot the SCPs, eQTL PIPs and GWAS PIPs for each individual SNP, respectively. No single SNP shows a particular high posterior probability in any of the three plots, but the cumulative regional probabilities from all the SNPs are all high. Panel D plots the pairwise LD pattern, measured by *r*^2^, for the 54 SNPs and indicates that all SNPs are tightly linked.

## Supplementary Figures

**Figure S1:**
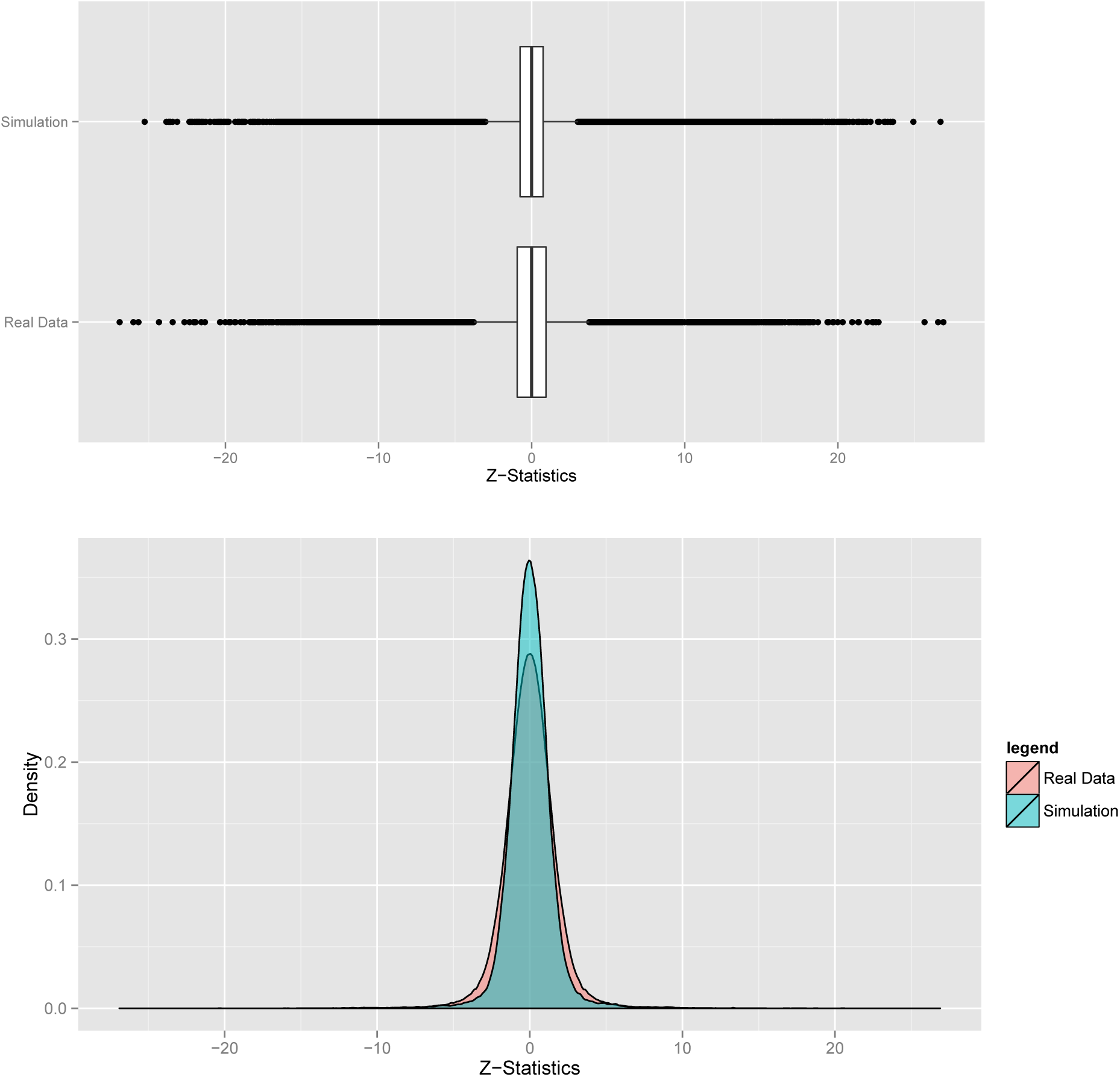
Comparison of the simulated summary *Z*-statistics and the observed data from height GWAS Wood *et al.* (2014). The overall distributional patterns of *z*-statistics are quite similar. The boxplot indicates that the extreme values from the two distributions are very much comparable; the density plot suggests the simulated *z*-statistics are more concentrated around 0, hence slightly conservative.

**Figure S2:**
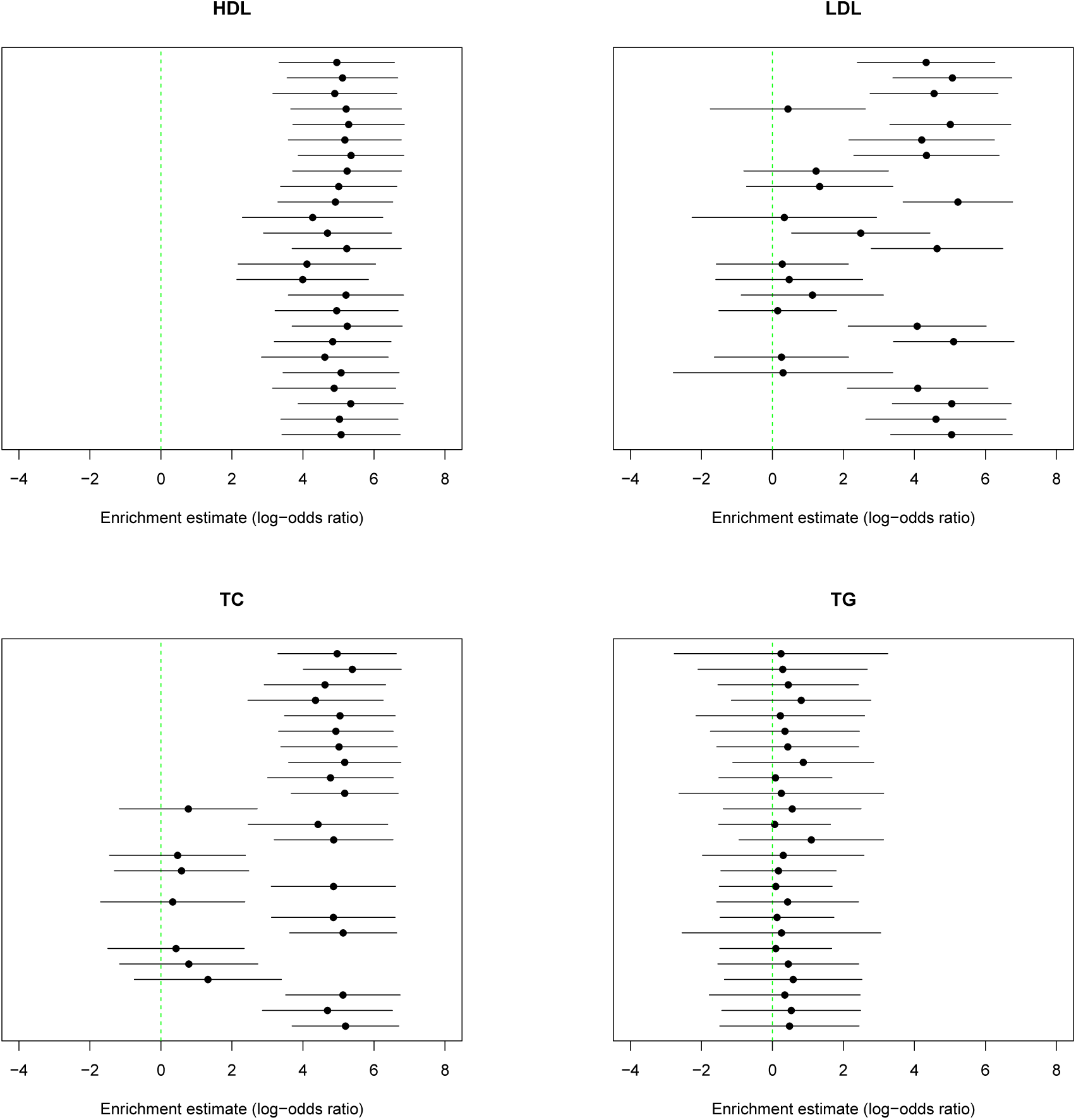
Individual estimates and their corresponding 95% confidence intervals from each imputed eQTL annotation data set in enrichment estimate of the four blood lipids traits.

